# The mesodermal source of fibronectin is required for heart morphogenesis and cardiac outflow tract elongation by regulating cell shape, polarity, and mechanotransduction in the second heart field

**DOI:** 10.1101/2022.10.28.514299

**Authors:** Cecilia Arriagada, Evan Lin, Michael Schonning, Sophie Astrof

## Abstract

Failure in the elongation of the cardiac outflow tract results in congenital heart disease due to ventricular septum defects and misalignment of the great vessels. The cardiac outflow tract lengthens via accretion of progenitors derived from the second heart field (SHF). SHF cells in the splanchnic mesoderm are exquisitely regionalized and organized into an epithelial-like layer forming the dorsal pericardial wall (DPW). Tissue tension, cell polarity, and proliferation within the DPW are important for the addition of SHF-derived cells to the heart and elongation of the cardiac outflow tract. However, the genes regulating these processes are not completely characterized. Using conditional mutagenesis in the mouse, we show that fibronectin (*Fn1)* synthesized by the SHF is a central regulator of epithelial architecture in the DPW. *Fn1* is enriched in the anterior DPW and mediates outflow tract elongation by balancing pro- and anti-adhesive cell-ECM interactions and regulating DPW cell shape, polarity, cohesion, proliferation, and mechanoresponsiveness. Our studies establish that Fn1 synthesized specifically by the mesoderm coordinates multiple cellular behaviors in the anterior DPW necessary for elongation of the cardiac outflow tract.

## Introduction

Congenital heart disease (CHD) is one of the most prevalent forms of birth defects, affecting approximately 40,000 newborns in the U.S. each year (Tsao et al., 2022). Therefore, it is important to understand the cellular and molecular mechanisms governing cardiac morphogenesis and alterations that cause CHD. At least three different sources of progenitors give rise to the heart (Zhang et al., 2021). Among these, progenitors from the second heart field (SHF) primarily contribute to the right ventricle, the outflow tract (OFT), and portions of the interventricular septum and the atria (Cai et al., 2003; Dyer and Kirby, 2009; Kelly et al., 2001; Kirby, 2007; Mjaatvedt et al., 2001; Verzi et al., 2005; Waldo et al., 2001). Correct formation and elongation of the OFT are necessary for the proper morphogenesis of the arterial pole of the heart, e.g., for intraventricular septation and correct positioning of the great vessels relative to the cardiac chambers (Cortes et al., 2018; Ward et al., 2005). Defects in these processes give rise to some of the most common and severe types of CHD (Tsao et al., 2022).

The OFT is a single tube connecting the heart with the systemic circulation. In the mouse, the OFT elongates between embryonic day (E) 8.5 and E10.75 of development by the addition of SHF cells to the nascent heart tube (Li et al., 2016; Meilhac and Buckingham, 2018; Sinha et al., 2012; Sun et al., 2007). The decreased contribution of SHF cells results in defective OFT elongation, leading to aberrant cardiac septation and defective alignment of the arterial pole of the heart with the ascending aorta and the pulmonary trunk (Waldo et al., 2005; Ward et al., 2005; Yelbuz et al., 2002).

SHF cells in the dorsal pericardial wall (DPW) (Figure S1A) directly behind the forming heart exhibit epithelial properties such as apicobasal polarity (Cortes et al., 2018; Francou et al., 2014). Aberrations in SHF cell polarity and cohesion in the DPW result in shortened OFT and CHD (Cortes et al., 2018; Li and Wang, 2018; Soh et al., 2014). For example, *Tbx1*, a critical transcription factor implicated in 22q11.2 deletion syndrome, orchestrates the elongation of the OFT by modulating multiple cellular properties of cardiac progenitors in the anterior DWP, such as cell polarity, organization of the actin cytoskeleton, and epithelial tension (Francou et al., 2017; Francou et al., 2014). Meanwhile, components of the non-canonical Wnt signaling pathway, such as Vangl2, Wnt5a, and Disheveled (Dvl), contribute to OFT elongation either directly, by functioning within the OFT (Vangl2) (Ramsbottom et al., 2014), or by governing the apicobasal polarity of SHF cells in the posterior DPW (Wnt5a, Dvl) (Li et al., 2016; Sinha et al., 2015; Sinha et al., 2012). The latter involves the modulation of actin cytoskeletal dynamics and cell intercalation behaviors in the posterior DPW that are crucial for the movement of DPW cells anteriorly to join the elongating OFT (Li et al., 2019; Li et al., 2016; Sinha et al., 2015; Sinha et al., 2012; van den Berg et al., 2009).

In addition to being polarized, SHF cells in the DPW are regionalized along the anterior-posterior and mediolateral axes (Bajolle et al., 2008; Kelly, 2023). For example, *Osr1, Hoxb1*, and *Wnt5a* are expressed in the posterior DPW, while *Tbx1* is expressed in the anterior (Stefanovic et al., 2020). This regionalization is important for OFT elongation and its subsequent morphogenesis (Li and Wang, 2018; Stefanovic et al., 2020).

In our previous studies, we found that cardiac OFT was short in embryos with global deletion of the extracellular matrix (ECM) glycoprotein *Fn1* (Mittal et al., 2013); however, these mutants exhibited a plethora of other morphogenetic defects, including defects in left-right asymmetry, the development of the neural crest, endoderm, paraxial mesoderm, and vasculature, as well as severe global growth defects, making it challenging to discern the precise functions of fibronectin, as well as where and when these functions were specifically important for heart development (George et al., 1997; George et al., 1993; Mittal et al., 2013; Pulina et al., 2014; Pulina et al., 2011; Villegas et al., 2013). A conditional mutagenesis approach indicated that the expression of Fn1 and its major receptor integrin α5β1 in the *Isl1* lineage played a role in the OFT elongation (Chen et al., 2015). However, the OFT elongation defects in the mutants lacking Fn1 or integrin α5 expression in the *Isl1* lineage were mild and not fully penetrant (Chen et al., 2015), posing challenges for mechanistic investigations. In addition, since *Isl1* is expressed in multiple pharyngeal lineages the endoderm, mesoderm, surface ectoderm, and some neural crest populations (Cai et al., 2003; Engleka et al., 2012), the use of the *Isl1^Cre/+^* line to delete Fn1 did not allow us to pinpoint the specific tissue source(s) of Fn1 crucial for the OFT development (Chen et al., 2015).

Prior to heart formation, *Fn1* mRNA is broadly expressed in the paraxial and lateral plate mesoderm of early (1-3 somite) murine embryos (Pulina et al., 2011). During the early stages of heart development (6-23 somites), *Fn1* mRNA continues to be expressed in the splanchnic mesoderm, while its expression in differentiated cardiomyocytes becomes undetectable (Mittal et al., 2010). We noticed that by ∼E9.5, *Fn1 mRNA* and protein were regionalized along the anterior-posterior axis of the DPW, and both Fn1 mRNA and protein were enriched in the anterior. These observations and previous studies pointing to cell-type-specific and cell-autonomous roles of Fn1 *in vivo* and *in vitro* (Cseh et al., 2010; Turner et al., 2017; Wang and Astrof, 2016) led us to hypothesize that the regionalized expression of *Fn1* in the DPW is important for cardiac development.

Mesodermal populations expressing *Fn1 mRNA* contain cardiac progenitors. Since the development of SHF-derived cardiac structures is particularly affected in global Fn1-null embryos and embryos with conditional deletion of Fn1 in the *Isl1* lineage (Chen et al., 2015; Mittal et al., 2013), we hypothesized that Fn1 synthesized by cardiac progenitors in the SHF was important for heart development. The half-life of Fn1 protein *in vivo* is approximately 21 hours (Deno et al., 1983; Sakai et al., 2001). Therefore, to downregulate Fn1 protein before the formation of SHF-derived cardiac segments, we used the *Mesp1^Cre/+^* knock-in strain in which Cre is expressed at E6.5 (Saga et al., 1996; Saga et al., 1999).

In our study, we demonstrate that mesodermal Fn1 plays a crucial role in coordinating multiple cellular processes involved in the accrual of SHF cells from the DPW during OFT elongation. The absence of mesodermal Fn1 resulted in the imbalance of Fn1 and Tenascin C (TnC) proteins directly beneath the DPW. TnC is an ECM glycoprotein known to interfere with cell binding to Fn1 and cause cell rounding with the concomitant loss of cell polarity, cell cohesion, and nuclear accumulation of YAP (Chiquet-Ehrismann et al., 1988; Midwood et al., 2004; Radwanska et al., 2017; Sun et al., 2018). Alterations in these properties in the anterior DPW ultimately disrupted the normal elongation of the cardiac OFT in mutants lacking the mesodermal source of Fn1.

## Results

### Mesodermal synthesis of Fn1 is required for cardiogenesis and embryonic viability

The expression of the transcription factor Mesp1 is detectable at E6.5 and marks the earliest known cardiac progenitors; descendants of Mesp1-expressing cells give rise to the anterior mesoderm, including mesodermally-derived cells of the heart (Saga et al., 1996; Saga et al., 1999). Therefore, we used the *Mesp1^Cre/+^*strain to conditionally ablate *Fn1* expression from heart precursors. Although Mendelian proportions of all genotypes were recovered at E9.5, we observed some Fn1^flox/-^; *Mesp1^Cre/+^* mutant embryos at E9.5 with mild cardiac defects (compare Figures S1B-B1 with Figures S1C-C1 and Figures S1D-D1). The heartbeat rate in controls and mutant embryos isolated at 20-26 somite stages was indistinguishable (Figure S1E), and no hemorrhage or edema was observed at this stage. However, only 50% of the expected number of Fn1^flox/-^; *Mesp1^Cre/+^*embryos were recovered at E10.75 (Figure S1F-G1). All the recovered mutants at E10.75 had prominent cardiac edema, defective hearts, and were degenerating (Figure S1G-G1); embryos from this stage were excluded from all the subsequent analyses. No Fn1^flox/-^; *Mesp1^Cre/+^* mutants were recovered after E13.5 days of gestation (Supplemental Table 1). These data indicated that the expression of *Fn1* in the Mesp1 lineage is required for heart development and embryonic viability.

### Mesodermal Fn1 regulates the elongation of the cardiac OFT

The shape of the heart in mild mutants at E9.5 and mutants surviving to E10.75 suggested that mesodermal Fn1 played a role in the development of the cardiac OFT (Figure S1D-D1, S1G-G1). To test this hypothesis, we first examined embryos at E8.5 (8-12 somites), when the OFT begins to form. At this stage of development, we did not find alterations in any cardiac structures, and the nascent OFTs in mutants and controls were of similar lengths (Figure S2A-C). The heart remained similar in shape in control and mutant embryos at 18-19 somites, and the OFT length was comparable among control and mutant genotypes at this stage (Figure S2D-G1, quantified in Figure S2J-K). However, at 20-25 somites, OFT length in mutant embryos was shorter than in controls: both the proximal and the distal portions of the OFT were affected and had fewer myocardial cells at this stage (Figure S2H-I1, S2L-O). The distal OFT (yellow segment in Figure S2H-I1) was more severely affected than the proximal segment (cyan in Figure S2H-I1), where the decrease in the cardiomyocyte number was not statistically significant (Figure S2N-O). In control embryos, the right and the left ventricles align in the plane perpendicular to the embryonic anterior-posterior axis (Figure S2P). In mutants with shortened OFT, this alignment is distorted, and the right ventricle is shifted anteriorly due to the decreased OFT length (Li et al., 2016). The alignment of the right ventricle was similarly distorted in Fn1^flox/-^; *Mesp1^Cre/+^*mutants (Figure. S2Q-R). Together, these data indicate that mesodermal Fn1 mediates cardiogenesis by regulating OFT elongation.

### Mesodermal Fn1 regulates the epithelial architecture of the anterior dorsal pericardial wall

Cardiac OFT elongates through the addition of cardiac precursors from the SHF in the first and second pharyngeal arches and from the splanchnic mesoderm comprising the DPW (Cortes et al., 2018; Li and Wang, 2018; Meilhac and Buckingham, 2018). The length of the OFT remained comparable among controls and mutants until the ∼19^th^ somite stage of development, suggesting that the addition of SHF cells from the pharyngeal arches was not affected by the deletion of the mesodermal source of Fn1 (Meilhac and Buckingham, 2018). To understand the causes of OFT elongation defects, we analyzed the cellular architecture of the DPW prior to the onset of OFT defects. For precision in comparing similar anatomical regions in controls and mutants, we performed our measurements within the medial region of the DPW, anatomically delimited by the lateral edges of the OFT, as shown in Figure S1A1. We began by analyzing embryos at E8.5, the time point that preceded the onset of OFT elongation defects by ∼24 hours. Although controls and Fn1^flox/-^; *Mesp1^Cre/+^* mutants were indistinguishable at this stage, at the cellular level, anterior DPW cells of E8.5 mutants were disorganized (Figure 1A-J). Instead of a single layer of epithelial-like cells, as seen in control Fn1^flox/+^; *Mesp1^Cre/+^* embryos, cells in the anterior DPW of Fn1^flox/-^; *Mesp1^Cre/+^* mutants were multilayered (compare Figure 1A-A1 with Figure 1D-D1, and 2D cellular traces in Figure 1B with 1E). In controls, anterior DPW cells appeared ellipsoid, regularly oriented along the dorsal-ventral axis, and neatly stacked along the length of the DPW (Figure 1A1, 1B, and 1C). Conversely, SHF cells in the anterior DPW of the mutants were more spherical (Figure 1D1, 1E, 1F). To quantify differences in cell shapes in 2D and 3D, we measured cell circularity, aspect ratio, ellipticity, and cell volume in controls and mutants (Figure 1G-J). Elongated cells have circularity and aspect ratios that deviate from 1, while spherical cells have circularity indexes close to 1. Differences in circularity indexes and the aspect ratio among SHF cells in the anterior DPW were statistically significant between controls and mutants (Figure 1G-H). To determine if 3D cell shape was also affected in the mutants, we segmented individual SHF cells in the anterior DPW and measured their ellipticity. Ellipticity was lower in Fn1^flox/-^; *Mesp1^Cre/+^* mutants than in controls, indicating that in the absence of mesodermal Fn1, SHF cells in the anterior DPW of the mutants were more spherical (Figure 1I). The volume of individual SHF cells was not affected in the mutants at this stage (Figure 1J). The aberrant epithelial architecture of the anterior DPW persisted at E9.5 (Figure 1K-S). At this time, defects in OFT elongation became detectable in the mutants (Figure S2H-H1, S2I-I1). In addition to altered cell shape, DPW cell volume was also decreased in E9.5 mutants (Figure 1T). Remarkably, in the posterior DPW cell shapes of control and mutant cells were comparable (Figure S3A-L). These observations suggested that mesodermal Fn1 regulated cell shape specifically in the anterior DPW (Movies 1-4 for more details and other optical sections).

**Figure 1.**
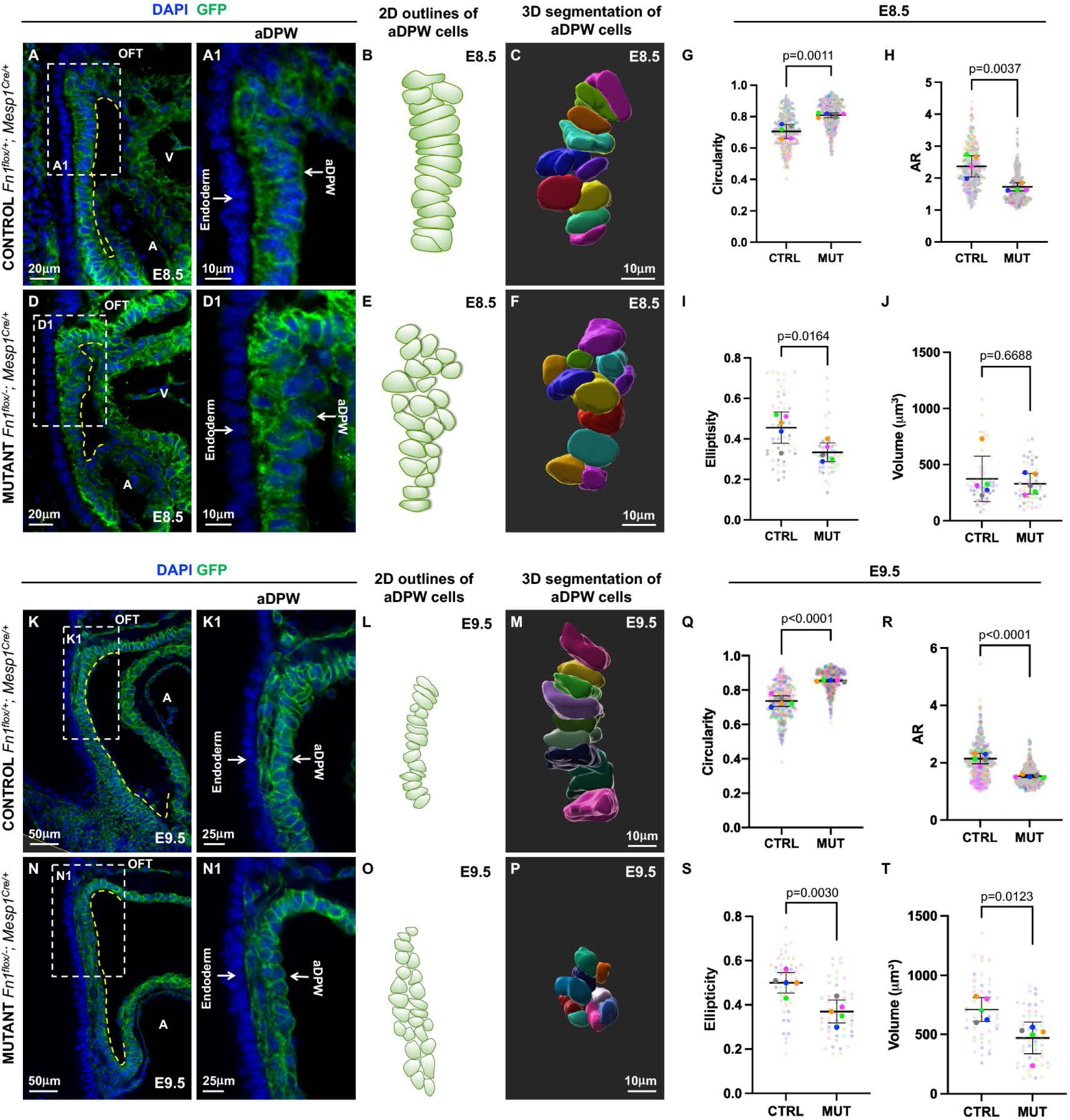
Mesodermal Fn1 regulates cell shape in the anterior DPW. **A-J.** E8.5 (9-11 s). **A-C.** Fn1^flox/+^; Mesp1^Cre/+^ (Control) and **D-F.** Fn1^flox/-^; Mesp1^Cre/+^ (Mutant) embryos stained to detect GFP (green) and nuclei (DAPI, blue). Regions marked by dashed rectangles in **A, D** are magnified to show the aDPW in **A1, D1. B, E.** Outlines of aDPW cells. **C, F.** 3D reconstructions of aDPW cells. **G-H.** 2D cell shape analyses. Small dots mark data points from each cell; large dots mark the average per embryo. Data from one embryo are marked with the same color. 320 cells from 5 control and 367 cells from 5 mutant embryos were analyzed. **G.** Cell circularity. **H.** Aspect ratio (AR). **I-J.** Ellipticity and volume of aDPW cells. 51 cells from 5 control and 50 cells from mutant embryos were analyzed. **K-T.** E9.5 (21-25s). **K-M.** Fn1^flox/+^; Mesp1^Cre/+^ (Control) and **N-P.** Fn1^flox/-^; Mesp1^Cre/+^ (Mutant) embryos, stained as above. Regions marked by dashed rectangles in **K, N** are magnified in **K1-N1. L, O.** 2D outlines of aDPW cells. **M, P.** 3D reconstructions of aDPW cells. **Q-R.** Cell shape analyses, color-coding is as above. 396 cells from 5 control and 436 cells from 5 mutants were analyzed. **Q.** Cell circularity. **R.** Aspect ratio. **S-T.** Ellipticity and volume of aDPW cells, n=50 cells from 5 control and 50 cells from 5 mutant embryos were analyzed. Means (horizontal bars) and standard deviations (error bars) are displayed; p values were calculated using 2-tailed, unpaired Student’s t-tests and the embryo averages. A-atrium, aDPW-anterior dorsal pericardial wall, OFT-outflow tract.

In summary, Fn1^flox/-^; *Mesp1^Cre/+^* mutant embryos had heart rates comparable with controls (Figure S1E) and lacked edema and hemorrhage at E9.5. Moreover, differences in 2D and 3D cell shapes observed at E8.5 preceded the onset of OFT shortening at E9.5, which was only very mild by the 20-26 somite stage (Figure S2H-I1, S2L-M). There were no differences in apoptosis between controls and mutants at this stage (Figure S4A-J). We also quantified the numbers of neural crest-derived cells in the pharyngeal region, since the absence of neural crest-derived cells could inhibit OFT elongation (Waldo et al., 2005), but we did not find differences in neural crest cell abundance at this stage (Figure S4K). Thus, the aberrant cellular architecture of the anterior DPW observed in embryos between E8.5-E9.5 (≤ 26 somites) is directly due to the deletion of *Fn1* in the mesoderm rather than any abnormalities in embryonic or cardiac morphology, cell survival, heartbeat, or hemodynamics.

### Mesodermal Fn1 regulates SHF cell orientation and apicobasal polarity in the anterior DPW

To determine the mechanisms by which mesodermal Fn1 regulates cell shape in the DPW, we asked whether the mesodermal source of Fn1 was important for apicobasal cell polarity. At E8.5, the Golgi apparatus is mainly positioned toward the apical surface of SHF cells along the entire length of the DPW (Figure 2A-A2, see Movie 5 for additional optical sections). This orderly positioning of the Golgi was reflected in the quantification of the angles of the Golgi position relative to the anterior-posterior axis (Figure 2A3, determined as shown in Figure 2C). In contrast, the orientation of the Golgi apparatus was disorganized in the anterior DPW in Fn1^flox/-^; *Mesp1^Cre/+^* mutants (Figure 2B-B3, quantified in 2D, see Movie 6). At E9.5, while the Golgi apparatus was consistently positioned apically in controls along the entire DPW (Figure 2E-E3, Movie 7), there was a striking disorganization of Golgi positioning in the anterior DPW in the mutants (Figure 2F-F3, Movie 8, quantified in 2G). However, in the posterior DPW, the Golgi apparatus was positioned toward the apical cell side in controls and mutants alike at E8.5 and E9.5 (Figure S3N-S). These experiments demonstrated that differences in cell shape and Golgi orientation in Fn1^flox/-^; *Mesp1^Cre/+^* mutants were specific to the anterior DPW.

**Figure 2.**
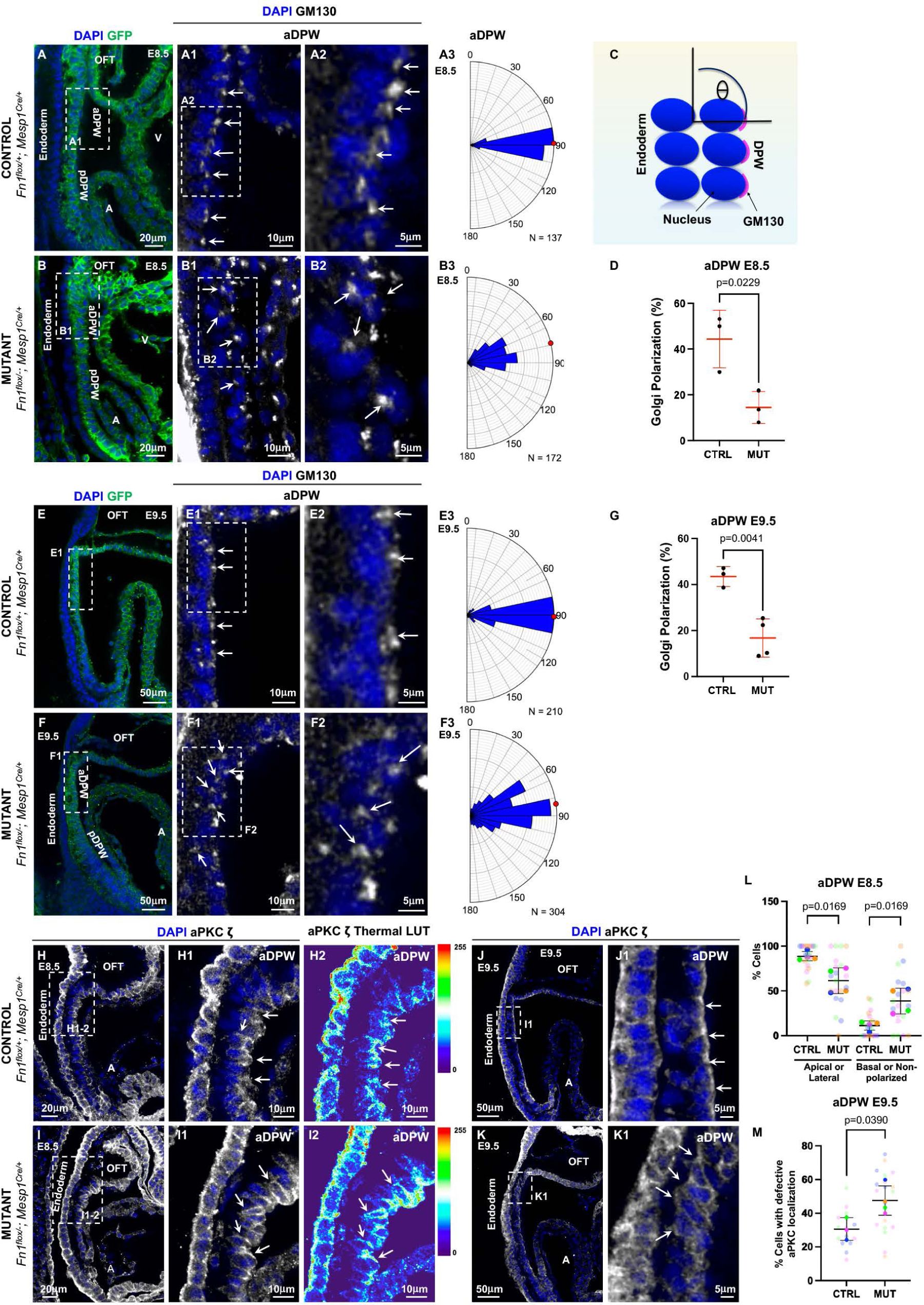
Mesodermal Fn1 regulates cell orientation and polarity in aDPW. **A-D.** E8.5 (9-11s). **A-A3.** Fn1^flox/+^; Mesp1^Cre/+^ (Control) and **B-B3.** Fn1^flox/-^; Mesp1^Cre/+^ (Mutant) embryos stained to detect GFP (green), GM130 (white), and nuclei (DAPI, blue). Boxes in **A, B** are magnified in the panels to the right. Arrows point to Golgi. **A3, B3.** Distribution of Golgi orientation angles in aDPW cells were measured as indicated in **C.** N=total number of cells analyzed in 3 controls and 3 mutants. **D.** Quantification of Golgi polarization. Golgi was considered polarized if the angle between the Golgi and the plane of the endoderm fell between 85° and 95°. **E-G.** E9.5 (20-22s). **E-E3.** Fn1^flox/+^; Mesp1^Cre/+^ (Control) and **F-F3.** Fn1^flox/-^; Mesp1^Cre/+^ (Mutant) embryos stained as above. Boxes in **E, F** are magnified in the panels to the right. Arrows point to Golgi. **E3, F3.** Distribution of Golgi orientation angles in aDPW cells measured as is indicated in **C.** N=total number of cells analyzed in 3 controls and 3 mutants. **G.** Quantification of Golgi polarization measured as in **D. H-K1.** Fn1^flox/+^; Mesp1^Cre/+^ (Control) and Fn1^flox/-^; Mesp1^Cre/+^ (Mutant) embryos were stained for aPKC **ζ** (white) and nuclei (DAPI, blue). Boxes mark regions magnified to the right. Arrows point to apical localization of aPKC **ζ** in controls and basolateral localization in mutants. **H-I2, L.** E8.5 (7-8s). **H2, I2.** aPKC **ζ** was color-coded according to fluorescence intensity, scales are on the right. **J-K1.** (20-22s), **L. E8.5. %** aDPW cells exhibiting a particular distribution of aPKC **ζ. M. E9.5** Quantification of cells in which aPKC was re-distributed away from the apical surface. DPW cells from 5 different slices were evaluated in 4 control and 4 mutant embryos (E8.5) or 3 controls and 4 mutants (E9.5). Each small dot marks data from one slice, and each large dot is an average of all slices per embryo. Data from the same embryo is marked by the same color. Means (horizontal lines) and standard deviations (error bars) are displayed; p values were calculated using 2-tailed, unpaired Student’s t-tests and the embryo averages. A-atrium, aDPW-anterior dorsal pericardial wall, OFT-outflow tract, pDPW-posterior dorsal pericardial wall, V-ventricle.

Like the disorganized Golgi orientation, apical cell surface markers were re-distributed in Fn1^flox/-^; *Mesp1^Cre/+^* mutants and were aberrantly localized at the basal surface of SHF cells in the anterior DPW at E8.5, the time before the onset of OFT elongation defects. These markers included podocalyxin (compare Figure S5A-A2 with Figure S5B-B2 quantified in Figure S5C, Movies 5-6) and atypical PKC ζ (aPKC ζ) (compare Figure 2H-H2 with Figure 2I-I2 quantified in Figure 2L, Movies 9-10). Changes in apical localization of aPKC ζ and podocalyxin in Fn1^flox/-^; *Mesp1^Cre/+^*mutants at E8.5 were specific for the anterior DPW, while in the posterior DPW, the localization of these proteins was not altered (Figure S5A3, B3, quantified in S5D, and Figure S5E-G). At E9.5, aPKC ζ localization remained disorganized in the aDPW in the mutants (compare Figure 2J-J1 with Figure 2K-K1, quantified in Figure 2M, Movies 11-12), but not in the posterior (Figure S5H-J); while the apically enriched localization of podocalyxin recovered at E9.5 (Figure S5K-M).

Cells in the anterior DPW are apically constricted and have a smaller apical surface area than cells in the posterior DPW (Francou et al., 2017). Consistent with these findings, we observed the enrichment of F-actin and an activated form of Myosin Light Chain 2 phosphorylated on Ser19 (pMLC), capable of driving the actomyosin contractility (Somlyo and Somlyo, 1994) at the apical cell surface of the aDPW (Figure 3A-B1, L-L2, N-N2). In contrast, not only were F-actin fibers disorganized and decreased in length in Fn1^flox/-^; *Mesp1^Cre/+^* mutants (Figure 3C-D1, E-K), but the apical enrichment of F-actin and pMLC was also lost (Figure M-M2, and 3O-O2). These changes in the organization of F-actin and contractile actomyosin cytoskeleton were specific to the anterior DPW (compare Figure 3 with Figure S6).

**Figure 3.**
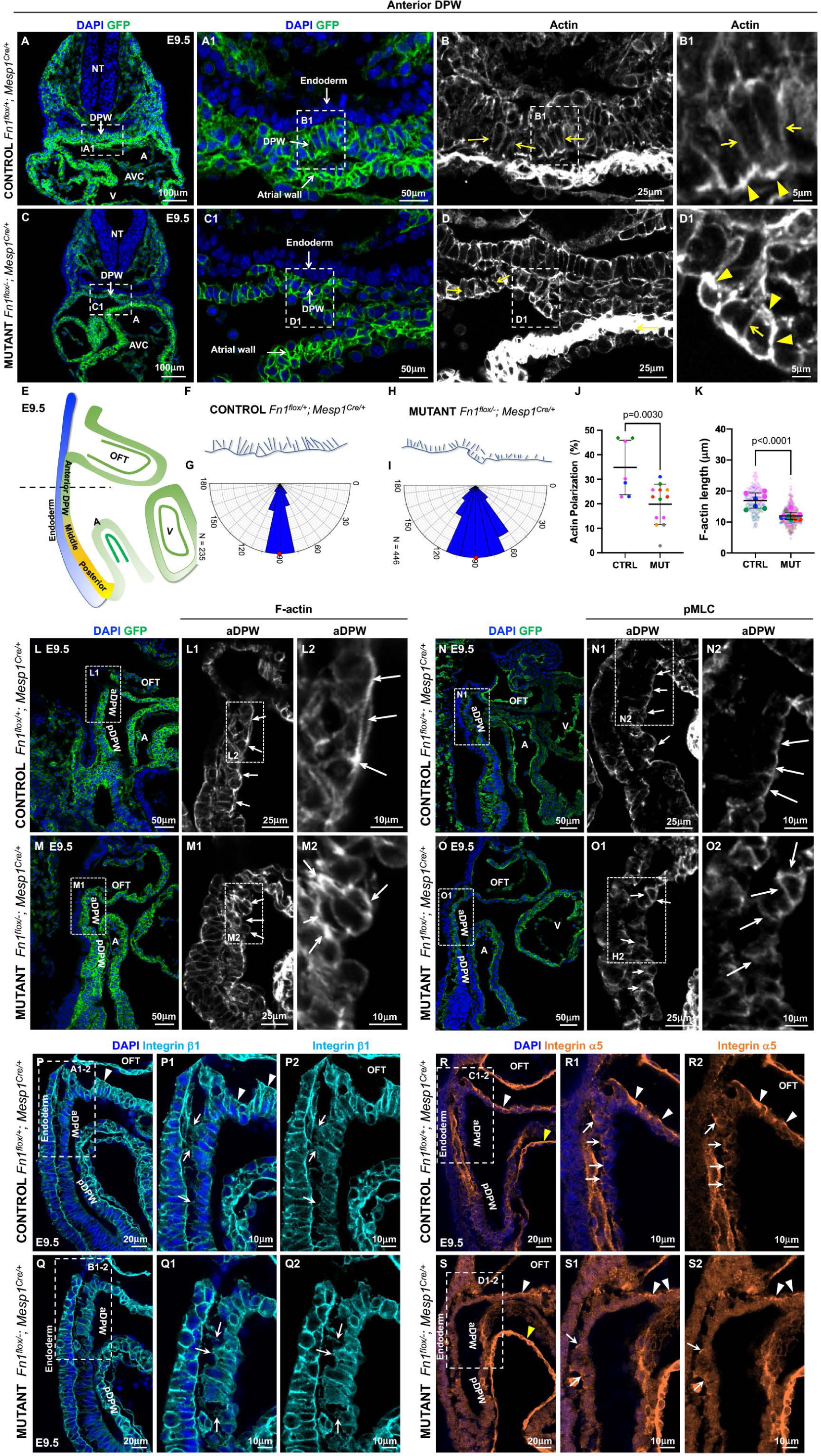
Mesodermal Fn1 regulates the actomyosin cytoskeleton and basolateral polarity in the aDPW. **A-B1.** Fn1^flox/+^; Mesp1^Cre/+^ (Control) and **C-D1.** Fn1^flox/-^; Mesp1^Cre/+^ (Mutant) embryos were dissected at E9.5 (21-26s). Transverse cryosections (dashed line in **E)** through the aDPW were stained to detect GFP (green), F-actin (white), and nuclei (DAPI, blue). Yellow arrows point to F-actin. Yellow arrowheads point to apically enriched F-actin in **B1** and re-distributed F-actin in **D1. E.** Schematic of different regions in the DPW. **F, H.** Traces of F-actin at apical and lateral cell borders. **G, I.** Angles made by lateral F-actin filaments (arrows in **B-B1, D-D1)** relative to cells’ apical surface. **J.** Quantification of actin polarization. An actin filament was considered polarized if was between 85° and 95° relative to the apical surface of the DPW. **K.** Quantification of the F-actin length at lateral borders (arrows in **B-B1, D-D1)** in the aDPW; 235 cells from 3 controls and 417 cells from 5 mutants were examined. Small dots mark the F-actin length of individual cells, and large dots mark their average length per slice. Dots from the same embryo are in the same color. Means (horizontal bars) and standard deviations (error bars) are displayed; p values were calculated using 2-tailed, unpaired Student’s t-tests and the embryo averages. **L-L2.** Fn1^flox/+^; Mesp1^Cre/+^ (Control) and **M-M2.** Fn1^flox/-^; Mesp1^Cre/+^ (Mutant) embryos were dissected at E9.5 (21-26s), and sagittal cryosections were stained to detect GFP (green), F-actin (white) and nuclei (DAPI, blue). Boxes in **L-M.** were expanded in **L1** and **M1** to show the aDPW. Boxes in **L1-M1** were expanded, in **L2** and **M2.** Arrows in **L1-L2** point to the apical enrichment of F-actin in the aDPW in controls. Arrows in **M1-M2** point to re-distributed F-actin in the aDPW of mutants. 6 controls and 3 mutants were analyzed. **N-N2.** Fn1^flox/+^; Mesp1^Cre/+^ (Control) and **O-O2.** Fn1^flox/-^; Mesp1^Cre/+^ (Mutant) embryos were dissected at E9.5 (21-26s) and sagittal cryosections were stained to detect GFP (green), pMLC (white) and nuclei (DAPI, blue). Boxes in **N-O** are expanded in **N1** and **O1.** Boxes in **N1-O1** are expanded in **N2** and **02.** Arrows in **N1-N2** indicate cells with apical localization of pMLC. Arrows in **01-02** show a defective basolateral localization of pMLC in mutants. 8 controls and 3 mutants were analyzed. **P-P2.** Fn1^flox/+^; Mesp1^Cre/+^ (Control) and **Q-Q2.** Fn1^flox/-^; Mesp1^Cre/+^ (Mutant) embryos were dissected at E9.5 (20-22s) and stained to detect Integrin β1 (turquoise) and nuclei (DAPI, blue). Regions marked by dashed rectangles in **P, Q** are magnified in **P1-P2, Q1Q2.** Arrows point to the basolateral localization of integrin β1 in aDPW cells in controls and mutants. **R-R2.** Fn1^flox/+^; Mesp1^Cre/+^ (Control) and **S-S2.** Fn1^flox/-^; Mesp1^Cre/+^ (Mutant) embryos were dissected at E9.5 (20-26s) and stained to detect integrin α5 (orange) and nuclei (DAPI, blue). Regions marked by dashed rectangles in **R, S** are magnified in **R1-R2, S1-S2.** Arrows point to the basally localized integrin α5 in aDPW cells in controls and mutants. Yellow arrowheads point to the basal localization of integrin α5 localization in the endocardium, white arrowheads point to the basal localization of integrin α in the OFT myocardium. aDPW-anterior dorsal pericardial wall, A-atrium, NT-Neural tube, OFT-outflow tract, V-ventricle.

To determine if the mesodermal Fn1 regulated the basal cell polarity, we assayed the localization of integrin subunits α5 and β1. Integrins are heterodimers of α and β subunits. 18 different α subunits and 8 distinct β subunits are known to form 24 different integrin heterodimers (Humphries et al., 2006). The integrin α5 subunit is only known to heterodimerize with integrin β1, while the integrin β1 subunit can heterodimerize with 12 different α subunits, which can form 12 different integrin heterodimers (Hynes, 2002). Integrin α5β1 is a major receptor for Fn1 (Humphries et al., 2006; Hynes et al., 1992; Yang et al., 1999; Yang et al., 1993).

In control DPW cells, the expression of integrin α5 and β1 subunits was enriched at the basolateral cell surface in the OFT (Figure 3P-P2, 3R-R2, white arrowheads), and the expression of integrin α5 subunit is also enriched basolaterally in the DPW (Figure 3R-R2, arrows). Although the localization of integrin β1 was not appreciably affected in the DPW (compare Figure 3P-P2 with 3Q-Q2, arrows), the localization of integrin α5 subunit at the basolateral surfaces was no longer prominent in Fn1^flox/-^; *Mesp1^Cre/+^* mutants (compare Figure 3R-R2 with Figure 3S-S2). Instead, the integrin α5 subunit was diffusely distributed in DPW cells (Figure 3S1-2) and was rarely enriched at their basal surface (Figure 3S2, arrows). As a control for integrin α5 staining, basal expression of integrin α5 subunit was evident in the OFT (Figure 3S1-2, white arrowheads), and in the endocardium (compare Figure 3S and 3R, yellow arrowheads) in the same optical section. Disrupted basolateral localization of integrin α5 subunit in the anterior DPW indicated that the basolateral cell polarity was disrupted in the aDPW cells in Fn1^flox/-^; *Mesp1^Cre/+^*mutants.

β1-containing integrin heterodimers collectively interact with many ECM proteins, including the ECM proteins underlying the DPW such as collagens, laminins, Tenascin C, and Fn1 (Alfano et al., 2019; Francou et al., 2014; Humphries et al., 2006; Hynes, 2002; Imanaka-Yoshida et al., 2003). In Fn1^flox/-^; *Mesp1^Cre/+^*mutants, β1-containing integrins are expressed at the basal surface of DPW cells, similar to controls (Figure 3P-P2, 3Q-Q2), allowing DPW cells to interact with ECM proteins underlying the DPW. Cell-ECM interactions result in the formation of cellular protrusions, and the number of cellular protrusions at the basal surface of DPW cells was not altered in Fn1^flox/-^; *Mesp1^Cre/+^* mutants (Figure S7A-C). In addition, integrin-mediated cell-ECM interactions can lead to the activation of Erk1/2 (Giancotti and Tarone, 2003), and the activation (phosphorylation) of Erk1/2 was not altered in Fn1^flox/-^; *Mesp1^Cre/+^* mutants (Figure S7D-I). Together with the re-distribution of F-actin and pMLC to the basal surface of DPW cells (Figure 3M2, O2), these observations suggest that in the absence of mesodermal Fn1, β1-containing integrin heterodimers on DPW cells interact with the underlying ECM and signal (Giancotti and Tarone, 2003; Miyamoto et al., 1996). However, the absence of enriched basal expression of the integrin α5 subunit in Fn1^flox/-^; *Mesp1^Cre/+^* mutants suggests that cell-ECM interactions mediated by β1-containing integrins *sans* α5β1 are inadequate for mediating proper cell shape and polarity in the anterior DPW.

In summary, our studies show that changes in cell shape, orientation, and polarity precede the changes in OFT length in Fn1^flox/-^; *Mesp1^Cre/+^* mutants and that altered cell shape and defective localization of GM130, aPKC ζ, F-actin, pMLC, and integrin α5 characterize the aberrant architecture of the anterior DPW of Fn1^flox/-^; *Mesp1^Cre/+^* mutants before and at the onset of OFT elongation defects. The observation that changes in cell shape, orientation, and polarity precede alterations in embryonic and heart morphology, and occur in the absence of hemorrhage or changes in heartbeat, indicates a direct involvement of mesodermal Fn1 in regulating the cellular architecture of the anterior DPW.

### Mesodermal Fn1 regulates mechanotransduction in the anterior dorsal pericardial wall

The transcriptional factor YAP is a central transducer of mechanical signals (Dupont, 2016). In response to changes in cell shape, integrin signaling, or mechanical tension, YAP translocates into the nucleus, regulating gene expression, actin cytoskeletal remodeling, cell migration, and proliferation (Borreguero-Munoz et al., 2019; Dupont et al., 2011; Dupont and Wickstrom, 2022; Elbediwy et al., 2016; Kim and Gumbiner, 2015; Wada et al., 2011). During heart development, YAP protein is diffusely localized in DPW cells at E8.5 (Francou et al., 2017) and (Figure S8A-D); but by E9.5, YAP translocates into the nucleus of DPW cells (Francou et al., 2017) and (Figure 4A-A2, nuclei are highlighted with dashed lines). Mechanical tension in the DPW at E9.5 is required for nuclear translocation of YAP, and YAP activity in the nucleus is important for DPW cell proliferation, the addition of the DPW cells to the OFT, and for OFT elongation (Francou et al., 2017).

**Figure 4.**
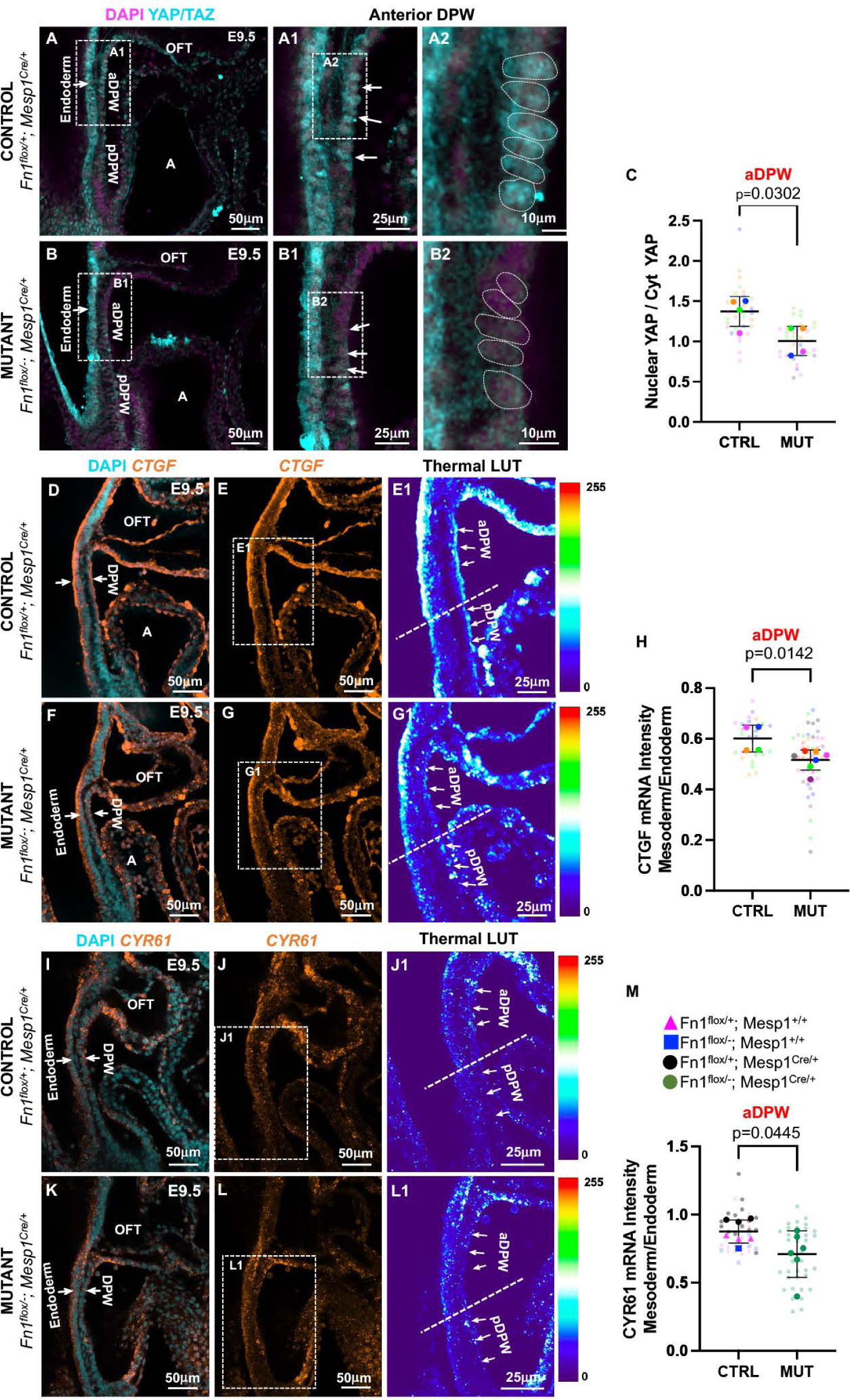
Mesodermal Fn1 regulates nuclear localization of YAP in the anterior DPW. **A-A2.** Fn1^flox/+^; Mesp1^Cre/+^ (Control) and **B-B2.** Fn1^flox/-^; Mesp1^Cre/+^ (Mutant) embryos were dissected at E9.5 (23-26s) and stained to detect nuclei (DAPI, magenta) and YAP protein (cyan). Nuclei in **A2, B2** are outlined by dashed lines. **C.** Quantification of the intensity ratio between nuclear and cytosolic YAP in the aDPW. 7-8 optical slices from 4 controls and 7-8 from 4 mutant embryos were analyzed. **D-E1.** Fn1^flox/+^; Mesp1^Cre/+^ (Control) and **F-G1.** Fn1^flox/-^; Mesp1^Cre/+^ (Mutant) embryos were dissected at E9.5 (23-26s) and labeled to detect *CTGF mRNA* (Orange) and nuclei (cyan). Rectangles in **E-G** are expanded in **E1-G1** and color-coded according to fluorescence intensity (scale is on the right). **H.** Quantification of *CTGF mRNA* fluorescence intensity in the aDPW (mesoderm) normalized to *CTGF mRNA* fluorescence intensity in the endoderm. n= 6 optical slices from 4 control and 9 mutant embryos were evaluated. Each small dot represents one slice, and each large dot is an average of all slices per embryo. Data from the same embryo are marked by the same color. **I-J1.** Fn1^flox/+^; Mesp1^Cre/+^ (Control) and **K-L1.** Fn1^flox/-^; Mesp1^Cre/+^ (Mutant) embryos were dissected at E9.5 (22-25s) and labeled to detect *CYR61 mRNA* (Orange) and nuclei (cyan). Rectangles in **J, L** are expanded in **J1, L1** and color-coded according to fluorescence intensity (scale is on the right). **M.** Quantification of *CYR61 mRNA* fluorescence in the aDPW normalized to the *CYR61 mRNA* fluorescence in the endoderm. n= 5-7 optical slices from 7 control and 6 mutant embryos were evaluated. **C, H, M.** Each small dot represents one slice, and each large dot is an average of all slices per embryo. Means (horizontal bars) and standard deviations (error bars) are displayed, 2-tailed, unpaired Student’s t - tests were used to determine p values using embryo averages. aDPW-anterior dorsal pericardial wall, OFT-outflow tract, pDPW-posterior dorsal pericardial wall.

Nuclear translocation of YAP is stringently regulated by cell shape and size (Aureille et al., 2019; Dupont et al., 2011; Elosegui-Artola et al., 2017), and this effect is dominant over stiffness (Cui et al., 2015; Dupont, 2016; Tang et al., 2013). Since DPW cells are rounder and smaller in Fn1^flox/-^; *Mesp1^Cre/+^* mutants than in controls at E9.5 (Figure 1T), we asked whether nuclear localization of YAP is altered in Fn1^flox/-^; *Mesp1^Cre/+^*mutants. For these experiments, we performed whole-mount immunofluorescence coupled with confocal imaging to detect YAP protein and analyzed YAP protein localization in the medial portion of the DPW, where cell shape, size, polarity, and actomyosin cytoskeleton were affected in Fn1^flox/-^; *Mesp1^Cre/+^* mutants (the region marked by vertical lines in Figure S1A1). We found that nuclear enrichment of YAP was significantly decreased in Fn1^flox/-^; *Mesp1^Cre/+^* mutants relative to controls in the anterior DPW (compare Figure 4A-A2 with Figure 4B-B2, quantified in Figure 4C), while in the posterior YAP localization was not affected (Figure S8E-G).

Downregulation of nuclear YAP is predicted to result in attenuated YAP-mediated transcription; therefore, we tested the expression of two direct transcriptional targets of YAP, *CTGF* and *Cyr61* (Stein et al., 2015). Consistent with our finding of diminished levels of nuclear YAP protein, *CTGF* and *Cyr61 mRNAs* were significantly downregulated in the anterior DPW of Fn1^flox/-^; *Mesp1^Cre/+^* mutants (Figure 4D-H and Figure 4I-M) but not in the posterior DPW (Figure S8H-I). The altered (more rounded) cell shape, re-distribution of the contractile actomyosin cytoskeleton from the apical surface basally, decreased cell volume, and decreased levels of nuclear YAP suggest an altered mechanical microenvironment (Perez Gonzalez et al., 2018) in the anterior DPW of Fn1^flox/-^; *Mesp1^Cre/+^*mutants.

### Regulation of proliferation by the mesodermal Fn1

A proliferation defect in the DPW (but not the heart or the endoderm) was already seen at E8.5 (Figure S9A-C, D-F, G-I), before the time when YAP becomes enriched in DPW nuclei (Figure S8A-D). At E9.5, proliferation was reduced in the cardiomyocytes of the OFT and RV (Figure S9J, K, L-M). Reduced proliferation was seen both in the anterior and posterior of the DPW in Fn1^flox/-^; *Mesp1^Cre/+^*mutants, but not in the adjacent endoderm (Figure S9N-O, Q-R, quantified in Figure S9T-U, X). These observations show that mesodermal Fn1 regulates the proliferation of DPW and its derivatives. The mechanism(s) by which the mesodermal Fn1 regulates DPW cell proliferation is likely independent of its role(s) in the regulation of cell shape, polarity, and YAP localization since decreased proliferation was seen in the posterior DPW where cellular architecture was not altered in the mutants and at E8.5, before the nuclear enrichment of YAP. However, the regulation of cell proliferation by Fn1 may be dependent on YAP in the anterior DPW at E9.5. The proliferation of myocardial cells in the left ventricle (LV), which are derived from the first heart field (FHF), was not affected in Fn1^flox/-^; *Mesp1^Cre/+^* mutants (Figure S9Y). This is consistent with the observation that the derivatives of the second heart field (SHF), the OFT and RV, are much more severely affected than the LV when Fn1 was deleted globally on C57BL/6J genetic background (Mittal et al., 2013). In summary, the proliferative defect in the DPW at E8.5 occurs along the entire length of the DPW before any changes in embryonic morphology and before the nuclear enrichment of YAP, indicating that the proliferation of DPW cells directly depends on the mesodermal expression of Fn1, and that altered proliferation in the DPW and the OFT contributes to defective OFT elongation in Fn1^flox/-^; *Mesp1^Cre/+^*mutants.

### Mesodermal Fn1 regulates cell cohesion in the SHF and collective cell migration

Cell-cell adhesion is important for collective cell migration (Friedl and Mayor, 2017). Adherens junctions form between the apical and basal cell surfaces and are important for mediating collective cell migration (Nelson, 2003; Weber et al., 2011; Weber et al., 2012). The adherens junction protein N-Cadherin is expressed in the DPW. Whereas N-Cadherin expression was not affected at E8.5 (Figure S10A-D2), N-cadherin levels were downregulated in the DPW of Fn1^flox/-^; *Mesp1^Cre/+^* mutants at E9.5 (Figure S10E-F2). Downregulation of N-Cadherin in cardiac progenitors within the anterior DPW causes their precocious differentiation (Soh et al., 2014). Consistent with these findings, we observed an expanded expression domain of the sarcomeric myosin heavy chain (assayed by the MF-20 antibody) in the anterior DPW of our mutants (Figure S10G-J). Alterations in cell shape, orientation, cell polarity, actin cytoskeleton, and YAP activity, together with the downregulation of N-Cadherin in the DPW, suggested that defective OFT elongation in Fn1^flox/-^; *Mesp1^Cre/+^*mutants could have resulted from aberrant collective cell migration of DPW cells to join the elongating OFT. To model the effect of cell-autonomous Fn1 deletion on cell migration, we obtained Fn1^flox/+^ and Fn1^flox/-^ mouse embryo fibroblasts (MEFs) and generated four (3 control and 1 Fn1-null) lines of MEFs (schematized in Figure S11A-C): The first control line consisted of Fn1^flox/+^ MEFs, wherein Fn1 protein can be expressed both from the floxed and the wild type *Fn1* alleles. In the second control group, Fn1^flox/-^ MEFs, Fn1 protein can be produced from the floxed *Fn1* allele, but not from the Fn1-null allele. To account for the potential effects of Cre recombinase and cell sorting on MEF behavior, a third control line, Fn1^Δ/+^ MEFs, was established. These MEFs were obtained through transient treatment of Fn1^flox/+^ MEFs with Cre recombinase, resulting in the recombined Fn1-null allele designated by Δ (Sakai et al., 2001). In Fn1^Δ/+^ MEFs, Fn1 is synthesized from the wild-type *Fn1* allele but not the Δ (null) allele. Finally, Fn1^Δ/-^ MEFs were generated by transient treatment of Fn1^flox/-^ MEFs with Cre recombinase; these MEFs are Fn1-null. All Cre-treated cells expressed GFP due to the presence of the ROSA^mTmG^ reporter allele and were sorted to obtain a population of GFP^+^ cells and the uninfected population of tdTomato+ cells (Figure S11C). Western blot analyses showed the downregulation of Fn1 protein expression in Fn1^Δ/+^, Fn1^flox/-^, and Fn1^Δ/-^ MEFs relative to Fn1^flox/+^ controls (Figure S11D-E).

Similar to the aDPW cells in Fn1^flox/-^; *Mesp1^Cre/+^*mutants, immunofluorescence and quantitative Western blot analyses showed that total and cell-surface levels of N-Cadherin were downregulated in Fn1^Δ/-^ MEFs (Figure S12A-G). N-Cadherin was also downregulated from MEF cell-cell adhesions (Figure S12D-D2). Since N-Cadherin was downregulated in Fn1^Δ/-^ MEFs, we used this model cell type and the monolayer scratch assay to determine whether the synthesis of Fn1 by MEFs was important for their collective cell migration. Scratch wounds made in Fn1^Δ/-^ MEF monolayers closed slower than in Fn1^flox/+^, Fn1^flox/-^, or Fn1^Δ/+^ MEF monolayers (Figure S12H-L). Consistent with the decrease in collective cell migration of Fn1-null MEFs during wound closure *in vitro*, we found that the number of myocardial cells in the distal OFT was decreased in Fn1^flox/-^; *Mesp1^Cre/+^* mutants (Figure S2N). Altogether, these studies suggest that mesodermal Fn1 regulates cell cohesion in the DPW at E9.5, facilitating the maintenance of the progenitor cell state in the anterior DPW cells and their collective movement to join the OFT, promoting its elongation.

### The mesodermal source of Fn1 is specifically important for the development of the anterior DPW and the OFT elongation

We next asked whether defects observed in the DPW of Fn1^flox/-^; *Mesp1^Cre/+^* mutants were due to decreased levels of Fn1 or were caused by the absence of Fn1 protein made specifically in the mesoderm. *Fn1 mRNA* and protein are dynamically and non-uniformly expressed in developing embryos (Chen et al., 2015; Mittal et al., 2010). Cells in the DPW and the pharyngeal endoderm are closely apposed in mid-gestation mouse embryos, and *Fn1* mRNA is synthesized by both tissues (Figure 5A-B2).

**Figure 5.**
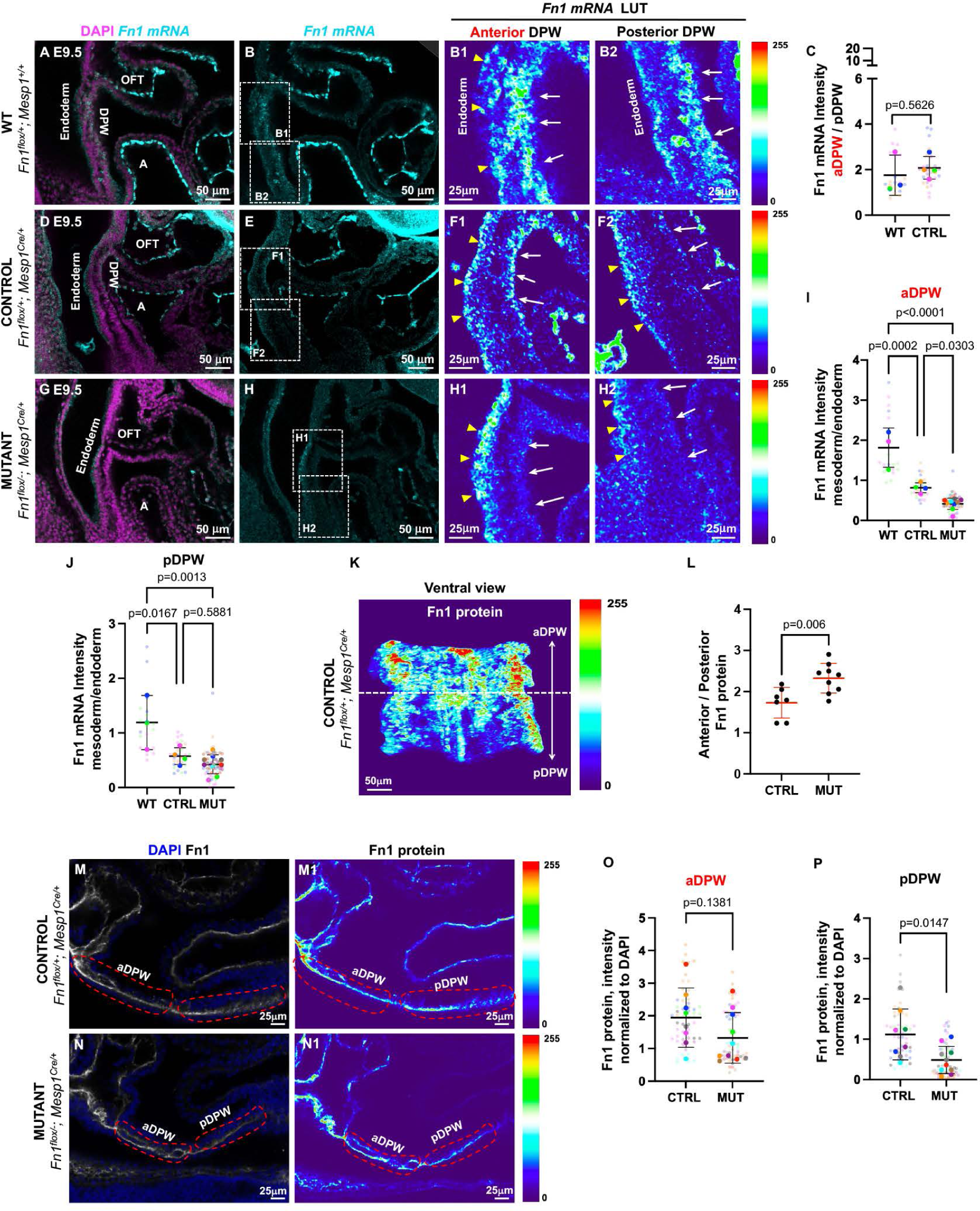
Expression patterns of *Fn1 mRNA* and protein in controls and mutants. **A-J.** Expression of *Fn1 mRNA.* Embryos were dissected at E9.5 (23-26s) and labeled with DAPI (magenta) and anti-sense Fn1 probes (turquoise). Regions outlined by the dotted rectangles in **B, E,** and **H** are expanded in panels to the right and color-coded according to fluorescence intensity (scales are on the right). White arrows point to DPW. Yellow arrowheads point to the endoderm. **A-B2.** Fn1^flox/+^; Mesp1^+/+^ (WT), **C.** *Fn1 mRNA* intensity ratio between aDPW and pDPW. *Fn1 mRNA* is enriched 2-fold in the aDPW relative to pDPW. **D-F2.** Fn1^flox/+^; Mesp1^Cre/+^ (Control). **G-H2.** Fn1^flox/-^·; Mesp1^Cre/+^ (Mutants). **I.** Normalized *Fn1 mRNA* intensity in the anterior and **(J)** posterior DPW, 6 different slices from 3 WT, 4 control, and 9 mutant embryos were analyzed. **K.** Ventral view of the DPW, Fn1 protein expression is color-coded according to fluorescence intensity. **L.** Ratio of Fn1 protein intensity shows Fn1 enrichment in the anterior DPW ot controls and mutants. **M, M1.** Fn1^flox/+^; Mesp1^Cre/+^ (Control) and **N, N1.** Fn1^flox/-^; Mesp1^Cre/+^ (Mutant) embryos were dissected at E9.5 (22-26s) and stained to detect Fn1 protein (white) and nuclei (DAPI, blue). **M1, N1.** Fn1 protein expression in the DPW is color-coded according to fluorescence intensity. **O-P.** Fn1 protein levels underlying the anterior DPW are unchanged in the mutants. Fn1 intensity in the aDPW and pDWP regions outlined by the red dash lines in **M1-N1** was normalized by DAPI intensity; 7 optical slices from 8 control and 10 mutant embryos were evaluated. Each small dot represents one slice, and each large dot is an average of all slices per embryo. Data from the same embryo are marked by the same color. Means (horizontal bars) and standard deviations (error bars) are displayed, 2-tailed, unpaired Student’s t-tests were used to determine p values using embryo averages. A-atrium, aDPW-anterior dorsal pericardial wall, LV-left ventricle OFT-outflow tract, pDPW-posterior dorsal pericardial wall.

At E9.5, *Fn1* mRNA was enriched nearly 2-fold in the anterior DPW relative to the posterior (Figure 5A-B2, quantified in Figure 5C). This enrichment is particularly evident in control Fn1^flox/+^; *Mesp1^Cre/+^*embryos (Figure 5D-F2). In these embryos, one allele of Fn1 is deleted in the mesoderm, resulting in a ∼50% decrease in *Fn1 mRNA* in the DPW relative to the wild type (Figure 5I). In Fn1^flox/-^; *Mesp1^Cre/+^* mutants, levels of *Fn1* mRNA in the entire DPW were low (Figure 5H-I, see panels H1-H2 for the anterior and posterior DPW, respectively; quantified in 5I-J). However, *Fn1 mRNA* in Fn1^flox/-^; *Mesp1^Cre/+^* mutants was still synthesized by the endoderm (Figure 5H1-H2, yellow arrowheads). The loss of *Fn1 mRNA* in the mesoderm but not in the endoderm is consistent with the known specificity of Cre expression in the *Mesp1^Cre/+^* strain (Saga et al., 1996; Saga et al., 1999)

Similar to *Fn1* mRNA, Fn1 protein was also enriched in the anterior DPW compared to the posterior at E9.5 (Figure 5K-L). Despite the efficient downregulation of *Fn1 mRNA* in the mesoderm of the DPW, the synthesis of *Fn1* mRNA by the endoderm (Figure 5H1, yellow arrowheads) resulted in a similar accumulation of Fn1 protein in the aDPW of controls and mutants (Figure 5M-M1, 5N-N1, quantified in Figure 5O). This is unlike the pDPW, where Fn1 protein was downregulated in Fn1^flox/-^; *Mesp1^Cre/+^*mutants (Figure 5P). Despite the presence of similar levels of Fn1 protein at E9.5, Fn1 derived from the endoderm did not rescue defects in the cellular architecture in the anterior DPW of Fn1^flox/^; *Mesp1^Cre/+^* mutants.

At E8.5, Fn1 protein is present in the DPW and the heart of controls (Figure S13A-B2) and is evenly distributed from the anterior to posterior DPW (Figure S13E), but it is downregulated in Fn1^flox/-^; *Mesp1^Cre/+^* mutants (Figure S13B1, D1, S13C-D1, and S13F-G). As a control, we measured Fn1 levels in the left ventricles, where Fn1 is synthesized by the endocardium (the mesoderm). Unlike in the ECM underlying the DPW, where Fn1 is derived both from the endoderm and the mesoderm, Fn1 protein levels were greatly diminished in the left ventricle of Fn1^flox/-^; *Mesp1^Cre/+^* mutants both at E8.5 and E9.5 (Figure S13B2, D2, H-M).

To determine the cell-type-specific functions of Fn1 in OFT elongation and the cellular architecture of the DPW and address the possibility that differences in Fn1 protein levels in the ECM underlying the DPW at E8.5 contributed to defective cellular architecture in the aDPW of Fn1^flox/-^; *Mesp1^Cre/+^*mutants, we deleted Fn1 using *Sox17^2A-iCre/+^* strain, in which Cre recombinase is expressed both in the endoderm and endothelium (Engert et al., 2009), Movies 13-14. In Fn1^flox/-^; *Sox17^2A-iCre/+^* mutants, *Fn1 mRNA* was downregulated in the endoderm, but it was still synthesized by the SHF cells of the DPW (compare Figure S14A-A1 with Figure S14C-C1). Although levels of Fn1 protein underlying the DPW were significantly reduced in Fn1^flox/-^; *Sox17^2A-iCre/+^* mutants (compare Figure S14B-B1 with Figure S14D-D1, quantified in S14E), the length of the cardiac OFTs were comparable between Fn1^flox/-^; *Sox17^2A-iCre/+^*mutants and all their genotypic controls (Figure S14F-I). The epithelial organization of the DPW and DPW cell shapes, assayed by detecting cell borders, were also not affected in Fn1^flox/-^; *Sox17^2A-iCre/+^* mutants (Figure S14K-N). Similarly, DPW cell orientation and polarity in Fn1^flox/-^; *Sox17^2A-iCre/+^* mutants were comparable with controls, and we did not see differences in the localization of GM130 or aPKC ζ (Figure S14O-V). These data indicate that synthesis of Fn1 specifically by the mesoderm is required for the proper cellular architecture of the DPW and the outflow tract elongation. Together, these studies establish a mesoderm-cell-autonomous role of Fn1 in the regulation of OFT development.

### Cell-autonomous requirement for Fn1 can be explained by the preferential integrin activation and Fn1 fibrillogenesis in cells synthesizing Fn1

Fn1 fibrillogenesis is mediated by active α5β1 integrins in the process wherein Fn1-α5β1 assemblies move along the cell surface from the cell periphery toward the cell center (Lu et al., 2020; Pankov et al., 2000; Tomer et al., 2022; Zamir et al., 2000). The centripetal movement of Fn1-α5β1 complexes is accompanied by the formation of long Fn1 fibrils on the extracellular side of the cell, mirrored by similarly elongated integrin α5β1-associated signaling assemblies on the cytoplasmic side (Lu et al., 2020; Pankov et al., 2000; Tomer et al., 2022; Zamir et al., 2000). The process of Fn1 fibrillogenesis is integrally linked to Fn1 signaling and is required for gastrulation, vascular development, and cartilage condensation; the blockade of Fn1 fibrillogenesis interferes with these processes (Chiang et al., 2009; Rozario et al., 2009; Singh and Schwarzbauer, 2014; Zhou et al., 2008).

The absence of distinct enrichment of integrin α5 at the basal surface of DPW cells in Fn1^flox/-^; *Mesp1^Cre/+^* mutants (Figure 3S-S2) suggested that DPW cells did not assemble fibrillar adhesions containing Fn1 and integrin α5β1, and that α5β1 integrins expressed by DPW cells were inactive (Clark et al., 2005). To test the hypothesis that the synthesis of Fn1 in cells expressing integrin α5β1 was important for the activation of α5β1 integrins, we generated Fn1^Δ/-^ mouse embryo fibroblasts (Fn1-null MEFs, Figure S11) to model the effects of cell-autonomous deletion of Fn1. Fn1 synthesis is not required for integrin trafficking to the cell surface (Cseh et al., 2010). Consistent with this, we found that integrin α5 was synthesized and transported to the cell surface in Fn1-null MEFs (Figure 6A-C). However, cell surface integrin α5 was not organized into fibrillar adhesions in Fn1-null MEFs compared with controls, even though Fn1 was present in the cell medium (compare Figure 6E1 with 6D1, arrows point to abundant fibrillar adhesions in Fn1-producing MEFs in Figure 6E1 and the rare fibrillar adhesions observed in Fn1-null MEFs in Figure 6D1).

**Figure 6.**
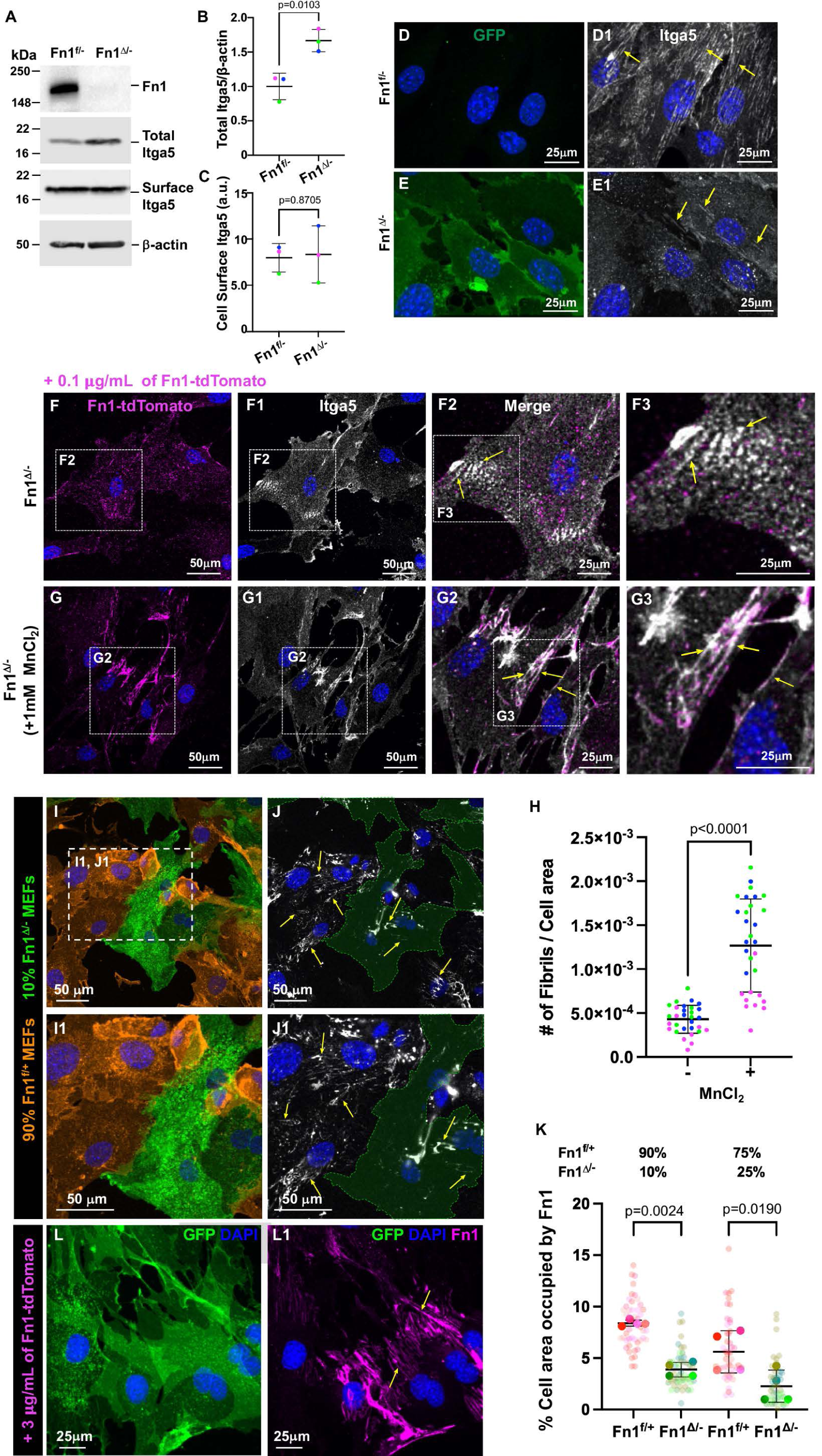
Cell-autonomous Fn1 regulates integrin α5 activation and Fn1 fibrillogenesis. **A.** Western blot analysis of total cell lysates and cell surface proteins. Positions of molecular weight markers are indicated on the left. **B-C.** Densitometrical quantification of immunoblot signals, n = 3 for each genotype. Each experiment is represented by a dot of a different color. Means (horizontal bars) and standard deviations (error bars) are displayed; a 2-tailed, unpaired Student’s t-test was used to determine p values. **D-E1.** Immunostaining to detect GFP (green), integrin α5 (Itga5, white), and nuclei (DAPI, blue) in Fn1^flox/-^ and Fn1^Δ/-^ (Fn1-null) MEFs. See **Fig. S11** for the explanation of genotypes. Arrows point to integrin α5 fibrillar adhesions in control Fn1^flox/-^ and Fn1^Δ/-^ (Fn1-null) MEFs. **F-F3, G-G3.** Fn1^Δ/-^ cells were cultured with 0.1 µg/mL of Fn1-tdTomato without **(F-F3)** or with **(G-G3)** 1mM MnCI_2_ for 6 hrs. After fixation, cells were stained to detect mCherry (Fn1-tdTomato, magenta), and integrin α5 (white). Arrows in **F2-F3** point to non-fibrillar Fn1 and integrin α5 adhesions in cells without MnCI_2_ treatment, and to fibrillar adhesions in the presence of MnCI_2_ in **(G2-G3). H.** Number of Fn1 fibrils per cell area. N=3 independent experiments, measurements from each experiment are represented by dots of the same color. Means (horizontal bars) and standard deviations (error bars) are displayed; a 2-tailed, unpaired Student’s t-test was used to determine the p value. **l-K. Fn1 is preferentially assembled in a cell-autonomous manner.** 1.5×10^4^ MEFs (generated as in **Fig. S11)** were plated in 24-well plates at the following proportions; 90% Fn1^f/+^ (td-Tomato^+^, orange) and 10% Fn1^Δ/-^ (Fn1-null, GFP^+^, green). After 48 hours of co-culture, cells were fixed with 4% PFA. Cells were stained to detect Fn1 without permeabilization (white). Panels I**-I1, J-J1** show GFP (green) and tdTomato (orange), and Fn1 staining (white). Regions marked by dashed rectangles in **I** are magnified in I**1** and **J1.** Green dashed lines outline areas occupied by GFP+ cells (Fn1^Δ/-^) in **J-J1.** Note that Fn1 fibrils (white) are mainly present within Fn1-expressing cells (orange in I**-I1),** arrows. **K.** Quantification of the percentage of cell area occupied by Fn1 staining in Fn1^f/+^ and Fn1^Δ/-^ MEFs in co-cultures. **L-L1.** Fn1-null cells (GFP^+^) assemble ectopic Fn1 into fibrils (arrows) when it’s provided in microgram quantities. Fn1^Δ/-^ MEFs were incubated with 3 µg/mL of Fn1-tdTomato fusion protein for 24 hrs. Immunofluorescence for GFP (green, Fn1-null cells) and mCherry (magenta, Fn1) was performed. Fn1 fibrils are in magenta in **L1,** arrows.

Integrin α5β1 is the only known integrin that facilitates the assembly of Fn1 into long fibrils (Sechler et al., 2001; van der Flier et al., 2010; Wennerberg et al., 1996). Upon binding Fn1, integrin α5β1 undergoes extensive conformational changes resulting in its activation and engagement with the actin cytoskeleton and cytoplasmic signaling effectors (Takagi et al., 2003). The paucity of α5β1^+^ fibrillar adhesions in Fn1-null MEFs suggested that cell surface α5β1 integrins were inactive in cells that did not synthesize Fn1. To shift cell surface integrins into their active conformation, control and Fn1-null cells were incubated with 1 mM Mn_2_Cl (Anderson et al., 2022; Masumoto and Hemler, 1993); 0.1 μg/ml of fluorescently labeled cellular Fn1 was added to the medium to visualize Fn1 fibrils. In the absence of Mn_2_Cl, Fn1-null MEFs rarely formed fibrils, and Fn1 and integrin α5β1 were mainly localized in dot-like adhesions (Figure 6F-F3). In contrast, incubation of Fn1-null MEFs with 1 mM Mn_2_Cl led to the robust formation of long fibrillar adhesions containing both Fn1 and α5β1 (Figure 6G-G3, quantified in H). These experiments demonstrated that Fn1-null cells can generate Fn1 fibrils upon integrin activation and suggested that the synthesis of Fn1 and integrins in the same cell facilitates integrin activation and Fn1 fibrillogenesis. In the absence of Fn1 synthesis, Fn1-null MEFs are deficient in their ability to generate long Fn1 fibrils, despite the presence of the ectopically added MEF-derived Fn1 in the medium.

Defective DPW architecture and OFT elongation in Fn1^flox/-^; *Mesp1^Cre/+^* mutants indicated that the presence of Fn1 protein synthesized by the endoderm was not sufficient to rescue DPW architecture and OFT elongation when the mesodermal Fn1 was absent. To model Fn1 assembly by Fn1-null cells situated in proximity with Fn1-secreting cells, we co-cultured Fn1^flox/+^ and Fn1^Δ/-^ (Fn1-null) MEFs in the same dish at different proportions. However, even when Fn1^flox/+^ and Fn1^Δ/-^ MEFs were cultured at the ratio of 9:1, Fn1 fibril assembly by Fn1^Δ/-^ MEFs was defective compared with Fn1^flox/+^ MEFs, even though Fn1^Δ/-^ MEFs were surrounded by Fn1^flox/+^ MEFs (Figure 6I-K, note the presence of Fn1 fibrils on the surface of control cells and the scarcity of Fn1 fibrils on the surfaces of the adjacent Fn1-null cells, quantified in Figure 6K). As a control, we incubated Fn1^Δ/-^ MEFs with fluorescently tagged Fn1-tdTomato (Tomer et al., 2022) at a concentration of 3 μg/ml. This experiment showed that Fn1^Δ/-^ MEFs are capable of assembling ectopically added Fn1 when it is supplied in large quantities (Figure 6L-L1), consistent with prior studies using microgram quantities of ectopically-added Fn1 (Cseh et al., 2010; Shi and Sottile, 2011); however, Fn1 fibril formation is significantly enhanced on the surfaces of cells that secrete it.

The absence of distinct integrin α5 localization at the basal surface of DPW cells Fn1^flox/-^; *Mesp1^Cre/+^* mutants (Figure 3S1-S2), deficiency in fibrillar adhesions (Figure 6E1), and decreased potency in Fn1 fibrillogenesis in Fn1-null MEFs (Figure 6I-K) suggested that DPW cells in Fn1^flox/-^; *Mesp1^Cre/+^*mutants were deficient in their ability to assemble Fn1 fibrils. To test this, we examined Fn1 ECM distribution between the endoderm and the DPW. For these experiments, we incubated whole embryos with antibodies recognizing Fn1 and imaged stained embryos using confocal microscopy. Compared with Fn1^flox/+^; *Mesp1^Cre/+^*controls and Fn1^flox/-^; *Sox17^2A-iCre/+^* endodermal mutants, Fn1 fibrillogenesis was disrupted at the basal surface of DPW cells in the mesodermal Fn1^flox/-^; *Mesp1^Cre/+^*mutants (Figures 7 and S15; compare Fn1 localization in Figure 7B1 with 7F1, and S15C1 with S15G1). To quantify these changes, we measured Fn1 intensity averaged through 20 μm-thick optical slices of the DPW, as shown in Figure S15A1, wherein the yellow rectangle identifies the region where average Fn1 intensity was quantified along the dorsoventral axis. There was a gradient of Fn1 distribution with the highest levels directly underlying the endoderm and the lowest levels underlying the DPW (Figure 7B-B1, quantified in Figure 7I, where the origin marks the basal side of the endoderm).

**Figure 7.**
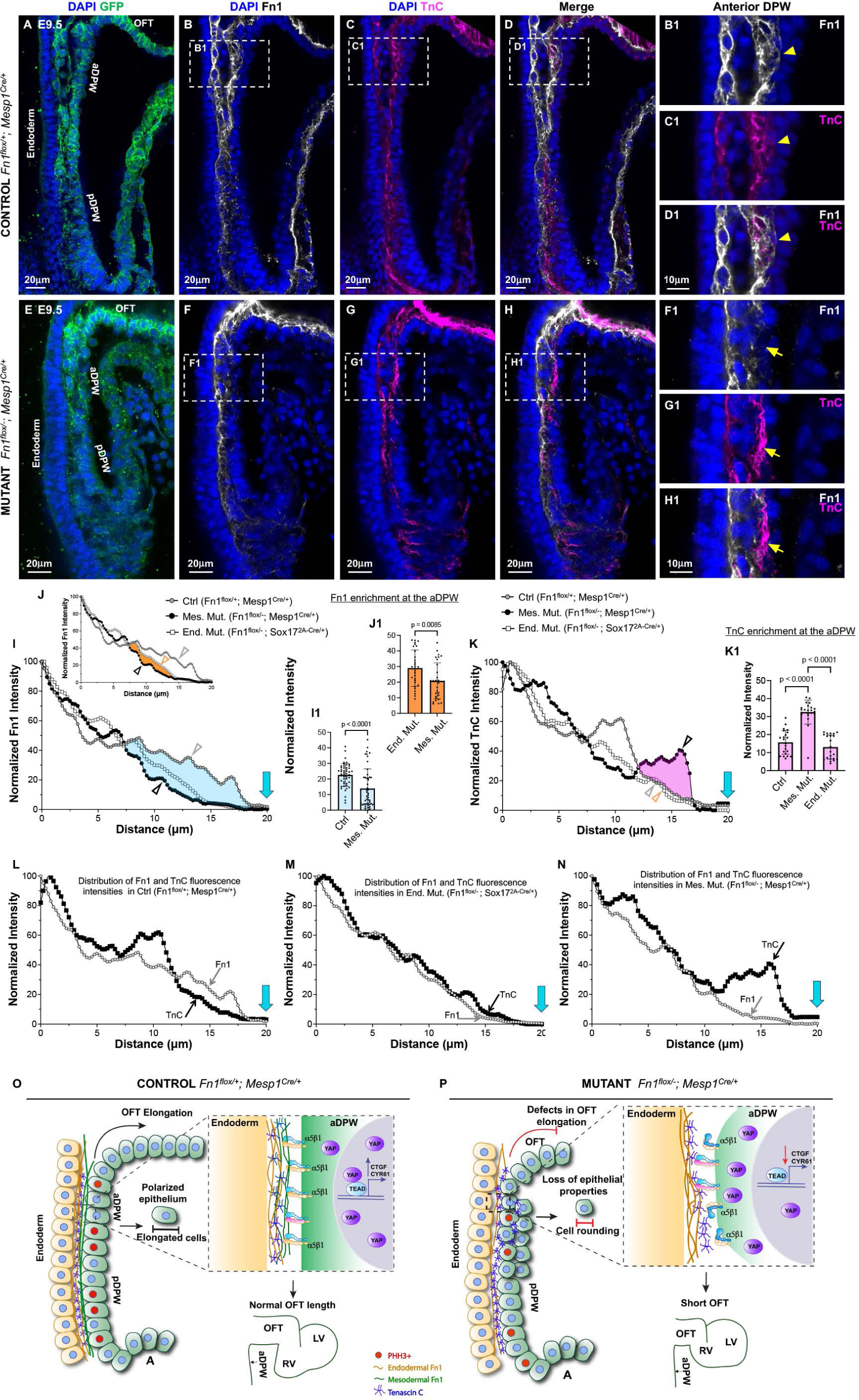
Mesodermal Fn1 regulates Tenascin C localization in the DPW. **A-D1.** Fn1^flox/+^; Mesp1^Cre/+^ (Control) and **E-H1.** Fn1^flox/-^; Mesp1^Cre/+^(Mutant) embryos were dissected at E9.5 (20-22s) and stained to detect GFP (green), Fn1 protein (white), TnC (magenta), and nuclei (DAPI, blue). Regions marked by dashed rectangles in **B-D** and **F-H** are magnified in **B1-D1** and **F1-H1.** Arrowheads point to Fn1 and TnC in controls, and arrows point to the altered distribution of Fn1 and TnC in mutants. **I-K1.** Distribution of Fn1 and TnC in the aDPW, distance from the basal surface of the endoderm is plotted on the x-axis at the origin. Blue arrows mark the apical edge of the DPW. **I-J.** Fn1 intensity profile in Fn1^flox/+^; Mesp1^Cre/+^ (Control, gray arrowhead), Fn1^flox/-^; Mesp1^Cre/+^(Mes. Mut., black arrowhead); and Fn1^flox/-^; Sox17^2A-iCre/+^ (End. Mut., orange arrowhead). The blue-shaded region in **I** is analyzed in I**1.** The orange-shaded region in **J** is analyzed in **J1. K-K1.** TnC intensity profile in Fn1^flox/+^; Mesp1^Cre/+^ (Control), Fn1^flox/-^; Mesp1^Cre/+^ (Mes. Mut.); and Fn1^flox/-^; Sox17^2A-iCre/+^ (End. Mut.) The pink-shaded region is analyzed in **K1**. **L-N.** Fn1 and TnC intensity profiles in each genotype. Note enriched Fn1 localization at the DPW in Control (**L**, gray arrow), **M.** Similar levels of Fn1 and TnC in Fn1^flox/-^; Sox17^2A-iCre/+^ (End. Mut.), but **N,** enriched TnC intensity at the DPW in the Fn1^flox/-^; Mesp1^Cre/+^ (Mes. Mut., black arrow). Blue arrows mark the apical edge of the DPW. **O-P. Model. O.** Fn1 synthesized by the mesoderm is important for integrin activation and Fn1 fibrillogenesis by aDPW cells. The mesodermal source of Fn1 mediates OFT elongation by regulating epithelial cell shape, orientation, polarity, proliferation, and mechanotransduction in the anterior DPW. **P.** In the absence of mesodermal Fn1, DPW-ECM interactions are altered, resulting in the loss of epithelial cell properties, decreased proliferation, and decreased cell volume, which lead to attenuated nuclear YAP localization and subsequent defects in the OFT elongation.

Fn1 intensity fell more abruptly in Fn1^flox/-^; *Mesp1^Cre/+^*mesodermal mutants than in Fn1^flox/+^; *Mesp1^Cre/+^* controls and Fn1^flox/-^; *Sox17^2A-iCre/+^* endodermal mutants (Figure 7I-J). The differences in the amount of Fn1 directly underlying the DPW between Fn1^flox/+^; *Mesp1^Cre/+^* controls and Fn1^flox/-^; *Mesp1^Cre/+^* mutants (region marked in blue, Figure 7I) were statistically significant (Figure 7I1). Similarly, the differences in Fn1 staining intensity between Fn1^flox/-^; *Sox17^2A-iCre/+^* endodermal and Fn1^flox/-^; *Mesp1^Cre /+^*mesodermal mutants (region marked in orange, Figure 7J) were also significant (Figure 7J1). Together, these experiments showed that Fn1 fibril assembly was decreased specifically at the basal surface of DPW cells in Fn1^flox/-^; *Mesp1^Cre/+^* mesodermal mutants relative to the endodermal mutants and controls.

### Mesodermal Fn1 is necessary to balance the adhesive and anti-adhesive properties of the ECM underlying the DPW

ECM contains regulatory proteins modulating cell-ECM interactions and, thereby, cell behaviors (Chiquet-Ehrismann and Tucker, 2011). Tenascin C (TnC) is one such modulatory protein present in the embryonic ECM (Imanaka-Yoshida et al., 2003). TnC modulates (counteracts) cell adhesion to Fn1 and modifies cell behavior, causing cell rounding, decreased cell-cell adhesion, and decreased proliferation of anchorage-dependent non-transformed cells (Chiquet-Ehrismann et al., 1988; Huang et al., 2001; Midwood et al., 2004; Orend et al., 2003; Radwanska et al., 2017). TnC binds the 13^th^ type III repeat of Fn1 within the HepII domain (Huang et al., 2001; Midwood and Schwarzbauer, 2002; Midwood et al., 2004; Orend et al., 2003) and blocks its interactions with Syndecan-4, an Fn1 co-receptor, which, along with integrin α5β1, is required for Fn1 fibrillogenesis and cell spreading (Galante and Schwarzbauer, 2007; Klass et al., 2000; Saoncella et al., 1999; Stepp et al., 2010). In endothelial cells, the modulatory effects of TnC depend on whether cells synthesize Fn1 (Radwanska et al., 2017). When TnC is incubated with endothelial cells producing Fn1, it enhances the formation of cell-cell junctions; however, the exposure of endothelial cells lacking Fn1 to TnC causes cell rounding and disruption of cell-cell adhesions (Radwanska et al., 2017).

TnC protein is expressed in the pharyngeal region underlying the DPW, the heart, and the cardiac outflow tract (Imanaka-Yoshida et al., 2003). We also detected TnC protein in the region between the DPW and the endoderm and noted that the pattern of TnC expression was similar to that of Fn1 (Figure 7C-D, C1-D1). To test the hypothesis that cell rounding and loss of DPW cell-cell junctions in Fn1^flox/-^; *Mesp1^Cre/+^* mutants could be due to the imbalance in Fn1 and TnC in the ECM underlying the DPW, we compared the distribution of Fn1 and TnC in control embryos with that in mutants in which Fn1 was deleted either in the mesoderm (Fn1^flox/-^; *Mesp1^Cre/+^*) or in the endoderm and endothelia (Fn1^flox/-^; *Sox17^2A-iCre/+^*). For these experiments, whole embryos were co-stained with antibodies to Fn1, TnC, and the lineage reporter GFP (Figure 7 and Figure S15). Although the total levels of TnC protein were unchanged in the ECM located between the DPW and the endoderm in Fn1^flox/-^; *Mesp1^Cre/+^* and Fn1^flox/-^; *Sox17^2A-iCre/+^* mutants relative to controls (Figure S15J-H), the localization of TnC protein at the basal surface of DPW cells was increased in Fn1^flox/-^; *Mesp1^Cre/+^* mutants (Figure 7G-H1, quantified in Figure 7K-K1). In controls and Fn1^flox/-^; *Sox17^2A-iCre/+^* endodermal mutants, Fn1 and TnC co-localized (Figure 7D, B1-D1, Figure S15B-E1, S15F-I1), and controls contained higher levels of Fn1 than TnC at the basal surface of the DPW (Figure 7L), while in the endodermal Fn1^flox/-^; *Sox17^2A-iCre/+^* mutants, Fn1 and TnC levels at the DPW were similar (Figure 7M). In contrast, TnC levels were greatly increased relative to Fn1 at the DPW of Fn1^flox/-^; *Mesp1^Cre/+^* mutants (Figure 7N). In addition, we observed regions underlying the DPW that contained TnC but lacked Fn1 in the mesodermal Fn1^flox/-^; *Mesp1^Cre/+^*mutants (Figure 7F1-H1). Quantitatively, there was a statistically significant enrichment of TnC at the aDPW in Fn1^flox/-^; *Mesp1^Cre/+^* mutants compared with the Fn1^flox/-^; *Sox17^2A-iCre/+^* endodermal mutants and Fn1^flox/+^; *Mesp1^Cre/+^* controls. In summary, these data suggest that the absence of Fn1 synthesis in the DPW leads to the enrichment of TnC at the basal surface of the DPW cells, underlying the defects in the DPW cellular architecture and OFT elongation.

Taken together, our studies suggest a model wherein Fn1 synthesized by the mesoderm engages and activates α5β1 integrins expressed by cardiac progenitors in the anterior DPW, facilitating Fn1 fibrillogenesis and signaling that regulates proliferation, cell shape, polarity, and cell volume in the DPW (Figure 7O). In the absence of mesodermal Fn1, DPW interactions with TnC underlying the DPW prevail (possibly through the binding to β1-containing integrins), resulting in cell rounding accompanied by the loss of cell polarity and cell-cell cohesion (Figure 7P). Both cell rounding and decreased cell volume in Fn1^flox/-^; *Mesp1^Cre/+^* mutants impede the nuclear translocation of YAP, leading to the loss of mechanoresponsiveness in the aDPW cells, and ultimately resulting in the defective elongation of the cardiac OFT (Figure 7P).

## Discussion

Here we show that Fn1 synthesized specifically by the mesoderm mediates the elongation of the cardiac OFT by regulating cell shape, polarity, orientation, proliferation, and the orderly stacking of cardiac progenitors in the anterior DPW. In Fn1^flox/-^; *Mesp1^Cre/+^* mutants, defects in these properties occur both before and at the onset of the OFT elongation defect, indicating that mesodermal Fn1 directly regulates cellular architecture and mechanotransduction in the anterior DPW.

The synthesis of *Fn1 mRNA* is regionalized in the SHF and enriched in the anterior DPW. Corresponding with the enrichment of *Fn1* mRNA, Fn1 protein is also locally enriched in the anterior DPW. This enrichment is a likely reason for the region-specific role of Fn1 in modulating cell shape, size, and polarity in the anterior DPW. Although *Fn1* synthesis was abolished in the mesoderm of Fn1^flox/-^; *Mesp1^Cre/+^* embryos, the levels of Fn1 protein between the anterior DPW and the endoderm in the mutants were not significantly different from controls at E9.5, likely due to the continued synthesis of *Fn1* mRNA by the adjacent endoderm. However, despite similar levels of Fn1 protein, Fn1 synthesized by the endoderm did not rescue defects in the cellular architecture of the anterior DPW in the mesodermal mutants. The deletion of Fn1 in the endoderm, which resulted in significantly reduced Fn1 levels, did not lead to defective DPW cell shape or polarity and did not cause defects in the OFT elongation. Together, these results indicate that Fn1 synthesized specifically by the mesoderm is necessary for regulating the epithelial properties in the anterior DPW and OFT elongation and that it does so in a cell-autonomous manner.

The conclusion above seemingly contrasts with the literature showing that Fn1 protein provided to cells in trans, either by adsorption to tissue culture plates or when added to cell supernatant, can be assembled into fibrils and signal (Avnur and Geiger, 1981; Miyamoto et al., 1996; Pankov et al., 2000; Peters et al., 1990; Shi and Sottile, 2011). Although we can also detect Fn1 fibrils when microgram amounts of Fn1 are supplied in the culture media of Fn1-null cells, experiments in this paper and others have shown that Fn1 fibrillogenesis is more efficient in cells synthesizing Fn1 (Cseh et al., 2010). In addition, we demonstrated that this effect can also be observed when Fn1-producing and Fn1-null cells are in direct contact with one another, or when they are in close proximity, as the DPW and endodermal cells *in vivo*.

Fn1 fibrillogenesis in Fn1-null cells can be rescued by either the addition of microgram quantities of Fn1 or by ectopically activating integrins, suggesting that co-expression of Fn1 and integrins in the same cell potentiates integrin activation and Fn1 fibrillogenesis. The lack of enriched expression of α5β1 integrins at the basal surface of DPW cells in Fn1^flox/-^; *Mesp1^Cre/+^* mutants and the decreased levels of Fn1 fibrils directly underlying the DPW cells is consistent with the notion that α5β1 integrins expressed by the DPW in Fn1^flox/-^; *Mesp1^Cre/+^*mutants are inactive, rendering DPW cells unable to bind Fn1 present in the ECM between the DPW and the endoderm. An alternative explanation for the deficiency in the ability of DPW cells to interact with Fn1 in the ECM of Fn1^flox/-^; *Mesp1^Cre/+^* mutants is the aberrant enrichment of TnC, which blocks cell binding to Fn1 (Huang et al., 2001; Midwood and Schwarzbauer, 2002; Midwood et al., 2004; Orend et al., 2003).

Our work and the work of others suggest that integrin α5β1 is the major transducer of Fn1 signals during embryogenesis (Chen et al., 2015; Mittal et al., 2013; Pulina et al., 2011; Takahashi et al., 2007; Wang and Astrof, 2016; Yang et al., 1999; Yang et al., 1993). Although multiple other integrins can bind Fn1, integrin α5β1 is the only known integrin that facilitates the production of long Fn1 fibrils (Sechler et al., 2001; van der Flier et al., 2010; Wennerberg et al., 1996). However, the deletion of integrin α5 in the Mesp1 lineage resulted in a much milder phenotype than the deletion of Fn1 (Liang et al., 2014). One explanation could be that the formation of short Fn1 fibrils by αv-containing integrins and/or by α4β1 integrins (van der Flier et al., 2010) partially compensates for the absence of α5β1 in the Mesp1 lineage. Alternatively, the potential co-transport of Fn1 and Fn1-binding integrins (α5β1, α4β1, or other β1- and αv-containing integrin heterodimers) in the same cell could be important to tilt the balance toward integrin interactions with Fn1 as opposed to other ECM proteins, like TnC. If this were the case, then in the absence of mesodermal Fn1, integrin interactions with TnC or other ECM proteins could prevail during their transport to the cell surface, attenuating their binding to the extracellular Fn1 remaining in the ECM of Fn1^flox/-^; *Mesp1^Cre/+^*mutants. If the co-transport of integrins with Fn1 were necessary for signaling by Fn1, then the deletion of any one of the Fn1-binding integrins would not be as severe as the deletion of Fn1.

Disruption of Fn1 fibrillogenesis in zebrafish and *Xenopus Laevis* embryos leads to defective planar cell polarity, cell shape, cadherin-mediated adhesion, and cell intercalation (Garavito-Aguilar et al., 2010; Kraft-Sheleg et al., 2016; Lackner et al., 2013; Marsden and DeSimone, 2001; Marsden and DeSimone, 2003; Rozario et al., 2009; Trinh and Stainier, 2004; Trinh et al., 2005). Conversely, disruption of cadherin-mediated adhesions in developing *Xenopus* embryos, interferes with Fn1 fibrillogenesis, resulting in tissue thickening and the shortening of the anterior-posterior axis due to defects in cell shape, intercellular intercalations, and convergent extension (Dzamba et al., 2009; Hirsh et al., 2018; Weber et al., 2011). Our data are consistent with these findings, indicating an indelible evolutionary-conserved role of Fn1 in the crosstalk between cell-cell and cell-ECM adhesions during tissue morphogenesis. In addition, our studies support a new paradigm that during morphogenesis, Fn1 regulates cell behaviors locally in a cell-autonomous manner and that Fn1 expressed by the anterior DPW is a central player coordinating multiple cellular behaviors regulating the development of the DPW and the elongation of the cardiac OFT. The deletion of the mesodermal source of Fn1 was accompanied by the aberrant localization of TnC. Thus, the resulting imbalance in the ECM composition could favor cell-TnC instead of cell-Fn1 interactions, leading to the observed phenotypes such as cell rounding, loss of cell polarity, and adherence junctions (Radwanska et al., 2017). In the future, it would be interesting to examine whether disruption of Fn1 in other model organisms results in aberrant localization of TnC.

Cell adhesion to Fn1 facilitates the remodeling of the actomyosin cytoskeleton and synergizes with growth factor signaling to promote the activation of Erk, the expression of cyclin D1, and cell cycle progression (Bohmer et al., 1996; Miyamoto et al., 1996; Roovers and Assoian, 2003; Walker et al., 2005). While the cellular architecture was mainly affected in the anterior DPW in Fn1^flox/-^; *Mesp1^Cre/+^*embryos, cell proliferation was affected along the entire length of the DPW as well as in the heart of E9.5 embryos, where the mesoderm is the sole source of Fn1 protein. Therefore, it is possible that in the DPW and the SHF-derived portions of the heart, Fn1 regulates proliferation via multiple mechanisms; in the anterior DPW, the defect in cell proliferation may be secondary to changes in cell shape, actomyosin cytoskeleton, and/or the nuclear localization of YAP, while in the posterior DPW and the heart, Fn1 may regulate proliferation directly by cooperating or coordinating signaling by growth factors known to be important for DPW proliferation, such as Fgf8 and Fgf10 (Dyer et al., 2010; Frank et al., 2002; Hutson et al., 2010; Hutson et al., 2006; Ilagan et al., 2006; Mittal et al., 2013; Park et al., 2008; Watanabe et al., 2010; Zhang et al., 2008).

The OFT and DPW are exquisitely regionalized, with distinct genetic programs regulating OFT elongation and the development of the arterial pole of the heart (Bajolle et al., 2006; Bajolle et al., 2008; Kelly, 2023; Li and Wang, 2018; Rammah et al., 2022; Ramsbottom et al., 2014; Stefanovic et al., 2020; Theveniau-Ruissy et al., 2008; Xia et al., 2019). Like Fn1, the expression of *Tbx1* – a major disease gene in 22q11.2 deletion syndrome – is enriched in the anterior DPW, where it regulates its cellular architecture (Francou et al., 2014; Jerome and Papaioannou, 2001; Lindsay et al., 1999; Lindsay et al., 2001; Merscher et al., 2001; Stefanovic et al., 2020). Deletion of *Tbx1* leads to defects in cell-extracellular matrix interactions, cell shape, cell organization, cell polarity, and actin cytoskeletal dynamics (Alfano et al., 2019; Francou et al., 2014), resulting in altered tension in the DPW and defective OFT elongation (Francou et al., 2017). Similar to *Tbx1*, *Fn1* mRNA synthesis and protein localization are enriched in the anterior DPW at E9.5, where Fn1 regulates cell shape, orientation, and biomechanical signaling. Similar phenotypes resulting from the deletion of *Tbx1* or *Fn1* in multiple contexts suggest the existence of genetic interaction between these pathways, a hypothesis worth testing in the future.

## Materials and Methods

### Antibodies

See Supplemental Table 2 for antibody information.

## Ethics Statement

All experiments involving vertebrate animals were approved by the Rutgers University Institutional Animal Care and Use Committee and were performed following federal guidelines for the humane care of animals.

### Mouse strains

C57BL/6J mice (cat # 0664) and *Rosa^mTmG/mTmG^* mice, *Gt(ROSA)26Sor^tm4(ACTB-tdTomato,-EGFP)Luo^* mice (Cat# 037456) (Muzumdar et al., 2007), were purchased from Jackson Laboratories. *Fn1^flox/flox^* mice were a gift from Reinhardt Fassler (Sakai et al., 2001). *Mesp1^Cre/+^* knock-in mice (Saga et al., 1996; Saga et al., 1999) were obtained from Dr. Yumiko Saga. *Fn1^+/-^*strain (George et al., 1997; George et al., 1993) was a gift from Richard Hynes. *Sox17^2A-iCre^* mice were a gift from Heicko Lickert (Engert et al., 2009). For experiments, *Fn1^+/-^* mice were crossed with *Mesp1^Cre/+^* knock-in mice to generate *Fn1^+/-^; Mesp1^Cre/+^* males which were propagated by mating with C57BL/6J females. To obtain embryos, *Fn1^+/-^; Mesp1^Cre/+^* males were crossed with *Fn1^flox/flox^; Rosa^mTmG/mTmG^* females and pregnant females were dissected at days specified in the Figures. To delete Fn1 in the endoderm and endothelium, Fn1^+/-^; Sox17^2A-iCre^ males were mated with *Fn1^flox/flox^; Rosa^mTmG/mTmG^* females and pregnant females were dissected at days specified in the Figures. Cre+ and Cre-negative controls were used in our studies, as specified in the figures and legends. Mice and embryos were genotyped according to published protocols (Engert et al., 2009; George et al., 1993; Saga et al., 1999). Embryos were staged by counting somites. The presence of the Y chromosome was determined by PCR using the primers using primers 5’-GCGCCCCATGAATGCATTTAT G-3’ and 5’-CCCTCCGATGAGGCTG-3’ to detect the sequence of the *SRY* gene. An equal number of male and female embryos were used.

### Cell culture and generation of cell lines

Mouse embryo fibroblasts (MEFs) were isolated from E13.5 embryos resulting from the mating of Fn1^+/-^ mice with Fn1^flox/flox^; ROSA^mTmG/mTmG^ mice. MEFs were cultured on plates pre-coated with 0.1% gelatin (Sigma, #G1890) dissolved in H_2_O and maintained in complete medium, which consisted of Dulbecco’s Modified Eagle’s medium (DMEM) with 4.5 g/L glucose, L-glutamine & sodium pyruvate (Corning, #10-013-CM) supplemented with 100 U/ml penicillin, 100 μg/ml streptomycin (Gibco, #15140-122), 1% v/v GlutaMax (L-glutamin, Gibco, #35050061) and 10% v/v fetal bovine serum (FBS, Gemini Biosciences, #100-106) in a humidified incubator with 5% CO_2_ at 37°C. Cells were tested for mycoplasma using the mycoplasma detection kit (ATCC, cat # 30-1012K) following the manufacturer’s instructions.

To generate MEFs lacking Fn1 (Fn1-null), or control, Fn1-heterozygous (Fn1-het) MEFs, Fn1^flox/+^; ROSA^mTmG/+^ and Fn1^flox/-^; ROSA^mTmG/+^ MEFs were infected with adenoviruses encoding Cre recombinase, Ad-Cre-IRED-GFP (Vector Biolabs, #1710). Cre-induced recombination not only leads to the ablation of Fn1 by recombining the floxed Fn1 sequence but also induces the expression of membrane-tethered GFP in these cells. For this experiment, 3×10^4^ MEFs were plated on gelatin-coated wells of 6-well plates. 24 hours later, cells were rinsed twice with Phosphate Buffered Saline 1X Sterile Solution (PBS, RPI Research Products, # P10400-500) and infected with Cre adenovirus diluted to the multiplicity of infection of 500 in DMEM without FBS. 20 hours post-infection, the medium was replaced by the complete medium. Although each of these cell types carries one ROSA^mTmG^ allele, only cells infected with Ad-Cre are constitutively and permanently marked by the expression of GFP due to the recombination of the reporter locus. GFP expression from the adenovirus is transient and dissipates after three days post-infection. After expansion, Ad-Cre-treated cells were sorted to generate Fn1^Δ/-^ (Fn1-null, GFP+), Fn1^flox/-^ (Fn1-heterozygous, GFP-), Fn1 ^Δ/+^ (Fn1-heterozygous, GFP+), and Fn1^flox/+^ (Fn1-wild type, GFP-) MEF lines. Western blot analysis was used to assay Fn1 expression.

### MEF protein lysates

Cells were plated in 6-well dishes at a density of 2×10^5^ cells per well. After 48 hours, cells were washed twice with cold PBS supplemented with 0.1 mM CaCl_2_ and 1 mM MgCl_2_ (PBS-Ca^2+^/Mg^2+^) and lysed with 400 μl of RIPA lysis buffer (50 mM Tris-Cl, 150 mM NaCl, 2 mM EDTA, 1% v/v NP-40, 0.5% w/v sodium deoxycholate, 0.1% w/v SDS, 1X protease inhibitor cocktail (Cell Signaling Technology, #5871), pH 8.0). The samples were centrifuged at 16,000 × g for 15 min at 4°C. Protein concentration was determined using BCA protein assay (Pierce™ BCA Protein Assay Kit, #23225). Then, lysis buffer was added to equalize protein concentration across samples. 4xNuPAGE™ LDS Sample buffer (Thermo Scientific, #NP0008) with 10% β-mercaptoethanol (Sigma, #63689) was added to each sample to the final concentration of 1X NuPAGE and 2.5% β-mercaptoethanol. Samples were heated to 95°C for 5 minutes prior to loading on gels.

### Immunoblotting and densitometry quantification

For immunoblotting, samples were resolved using 4-12% Tris-Glycine gel (Invitrogen, #XP04120BOX) at 130 V in 1x Tris-Glycine SDS Running Buffer (Invitrogen, # LC2675) for 1 hour 30 minutes. Proteins were then transferred to nitrocellulose membranes with a 0.2 µm pore size (Bio-Rad, # 1620122) and 1x Tris-Glycine, 15% methanol transfer buffer (10x Tris-Glycine buffer: 30g Tris Base, 144 g Glycine in 1L of ddH20, pH: 8.5). The transfer was carried out at 90 V for 2 hours in ice.

After the transfer, the nitrocellulose membranes were blocked with Intercept Blocking Buffer (Li-Cor, # 927-60001) for 1 hour. Primary antibodies were diluted in Intercept Antibody Diluent (Li-Cor, # 927-65001) and incubated overnight at 4°C. After 24h, the nitrocellulose membranes were washed three times with PBS containing 0.1% Tween for 10 min. We used IRDye 680RD Donkey anti-Rabbit IgG (#926-68073), and IRDye 800CW Donkey anti-Mouse IgG Secondary Antibody (#926-32212) as secondary antibodies in a 1:15000 dilution in Intercept Antibody Diluent (Li-Cor, # 927-65001). The nitrocellulose membranes were incubated with the secondary antibodies for 1 hour at room temperature. Finally, membranes were washed three times with PBS 0.1% Tween for 10 minutes and imaged using Li-Cor Odyssey 9120 Gel Imaging System (#ODY-2425). The density of unsaturated pixels was determined using ImageJ software (version 2.1.0/1.53c4). For each condition, protein bands were quantified from at least three independent experiments.

### Cell surface biotinylation

3.5×10^5^ cells were plated on gelatin-coated 10 cm plates. After 48 hours of growth, cells were washed twice with ice-cold PBS supplemented with 0.1 mM CaCl_2_ and 1 mM MgCl_2_ (PBS-Ca^2+^/Mg^2+^), followed by incubation with 1 mM EZ-Link Sulfo-NHS-LC-Biotin (Thermo Scientific, #21335) in PBS-Ca^2+^/Mg^2+^ for 30 min at 4°C. The biotinylation reaction was quenched by incubating cells with Tris-buffered solution (50 mM Tris-HCl, pH 7.4) for 10 min at 4°C. After two subsequent washes with PBS-Ca^2+^/Mg^2+^, cells were lysed with 600 μl of RIPA lysis buffer (50 mM Tris-Cl, 150 mM NaCl, 2 mM EDTA, 1% v/v NP-40, 0.5% w/v sodium deoxycholate, 0.1% w/v SDS, 1X protease inhibitor cocktail (Cell Signaling Technology, #5871), pH 8.0). To remove insoluble material, cell extracts were clarified by centrifugation at 16,000 x g for 15 min at 4°C. Soluble biotinylated protein concentration was determined using BCA protein assay (Pierce™ BCA Protein Assay Kit, #23225). 300 µg of protein per sample was incubated with 40 µl of Neutravidin-Agarose beads (Thermo Scientific, # 29202) pre-washed with RIPA lysis buffer. After 1 hour of incubation at 4°C, the Neutravidin-Agarose beads were washed 3 times with RIPA lysis buffer containing 1X protease inhibitor cocktail. 120 µl of 1X NuPAGE™ LDS Sample buffer (Thermo Scientific, #NP0008) with 2.5% β-mercaptoethanol (Sigma, #63689) and 1X protease inhibitor cocktail was added to beads and resolved by SDS-PAGE (Arriagada et al., 2020).

### Immunofluorescence microscopy

2×10^4^ cells were plated on gelatin-coated #1.5 round glass coverslips (Fisher Scientific, t#12-545-81) in 24-well dishes. After 48 hours, cells were washed with PBS at room temperature (rt) and fixed with 4% PFA (Fisher Scientific # 50-980-487, diluted in PBS) for 20 minutes at rt. Then, cells were washed three times with PBS for 5 minutes each and then used with or without permeabilization with PBS containing 0.1% Triton-X 100 (PBST) for 15 minutes. Cells were blocked with 5% Donkey serum (Sigma, #d9663) in PBST (blocking solution) for 30 minutes. After blocking, cells were incubated with the primary antibodies diluted in blocking solution overnight at 4°C, at dilutions indicated in Supplemental Table 2. Following 3 washes in PBST for 10 minutes each, cells were incubated with secondary antibodies diluted in blocking solution for 60 minutes at room temperature. Finally, cells were washed three times with PBST for 10 minutes, and then mounted using a 50% v/v Glycerol/Methanol solution.

### Co-culture of Fn1^Δ/-^ and Fn1^f/+^ MEFs and quantification of Fn1 incorporation into cellular ECM

A total of 1.5×10^4^ cells were plated in 24-well plates on coverslips coated with 1% gelatin solution, with the following proportions: 90% Fn1^f/+^ and 10% Fn1^Δ/-^ MEFs, or 75% Fn1^f/+^ and 25% Fn1^Δ/-^ MEFs in DMEM containing 10% FBS. After 48 hours, the medium was collected, and cells were fixed with 4% PFA 15 min at room temp. Cells were then washed with PBS and stained to detect Fn1 without permeabilization using anti-Fn1 primary antibodies and Alexa647-conjugated 2° antibodies. Antibody dilutions are listed in the Supplemental Table 2. Cells were mounted using Vectashield antifade mounting medium (Vector Laboratories, H-1000), GFP and tdTomato were visualized by imaging their native fluorescence. Fiji was used to segment and mask GFP^+^ (Fn1^Δ/-^ MEFs) or tdTomato^+^ (Fn1^f/+^) cells and to quantify the percentage of the masked areas occupied by Fn1+ pixels.

### Fn1^Δ/-^ incubation with cellular Fn1

1.5×10^4^ Fn1^Δ/-^ MEFs were plated in 24-well plates on coverslips coated with 1% gelatin. After 24 hrs, cells were incubated with 0.1 mg/mL of Fn1-tdTomato in DMEM containing 10% FBS for 24 hrs; This concentration of Fn1 was chosen after measuring Fn1 concentration in the medium of 90% Fn1^f/+^ and 10% Fn1^Δ/-^ MEF cells using the Mouse Fibronectin ELISA Kit (ab210967), which was 0.1 mg/ml. To treat cells with MnCl_2_, 1 M MnCl_2_ solution (Sigma, M1782) was diluted 1:100 in DMEM cell culture medium and then added to cells to a final concentration of 1 mM for 6 hrs. Cells were then fixed with 4% PFA for immunofluorescence with permeabilization to detect mCherry (Fn1-tdTomato), GFP, and integrin α5. The number of Fn1 fibrils was manually counted in each cell and normalized by total cell area.

### Monolayer Scratch Wound Assay

1×10^4^ cells were plated on gelatin-coated 8-well ibidi dishes (Ibidi, #80827) for 72 hours. When cells were confluent, the monolayer was scratched with a sterile 10 µl pipette tip. Cell debris was washed out with PBS, and incubated with imaging medium 100 μl FluoroBrite DMEM (Thermo Fisher Scientific, #A1896701) supplemented with 100 U/ml penicillin, 100 μg/ml streptomycin (Gibco, #15140-122), 1% v/v GlutaMax (L-glutamin, Gibco, #35050061) and 2% v/v fetal bovine serum (FBS, Gemini Biosciences, #100-106). Images were acquired right after performing the scratch and then after 14 hours. % of wound closure was measured as follows: % of Wound closure = *100*(A_t=0_ - A_t=14_)/ A_t=0_)*, *where A_t=0_ is the cell-free area at t=0 and A_t=14_ is the cell-free area at t=14 hrs*.

### Whole Mount Immunofluorescence Staining

Timed matings were set up to obtain embryos at specific embryonic stages. Vaginal plugs were checked daily, and noon of the day the plug was found was an embryonic day (E) 0.5 of gestation. Embryo dissections were aided using a Zeiss Stemi 2000C stereomicroscope. Embryos were dissected into ice-cold PBS; yolk sacs were kept for genotyping. After dissection, embryos were fixed with 4% PFA overnight at 4°C. Then, embryos were washed two times with ice-cold PBS and permeabilized with PBS containing 0.1% Triton-X 100 (PBST) overnight at 4°C. Embryos were blocked with 10% Donkey serum in PBST (blocking solution) overnight at 4°C. After blocking, embryos were incubated with the primary antibodies diluted in blocking solution for 3 days at 4°C. Following four 1 hour-washes with PBST, embryos were incubated with secondary antibodies diluted in blocking solution for 3 days at 4°C. Finally, after four washes with PBST for 1 hour each, embryos were embedded in 1% agarose (Bio-Rad Laboratories, #1613101), and cleared using methanol and Benzyl Alcohol (Sigma, # B-1042)/ Benzyl Benzoate (Sigma, #B-6630) as described (Ramirez and Astrof, 2020). For imaging, embryos were a placed between two #1.5 coverslips (VWR, #16004-312) separated by a rubber spacer (Grace Bio Labs, # 664113).

### Whole Mount Immunofluorescence in situ hybridization

After dissection, embryos were fixed with 4% PFA overnight at 4°C and washed twice with PBS. *In situ hybridization* assay carried out using the RNAscope® Multiplex Fluorescent v2 reagents (Advanced Cell Diagnostics, # 323110) as previously described (Nomaru et al., 2021). We used a C1 probe for Fn1 (#408181).

### Embryo Cryosection and Immunofluorescence Staining

After dissection, embryos were fixed with 4% PFA overnight at 4°C and washed twice with PBS. Embryos were then incubated with 10% sucrose in PBS for 4 hours at room temperature and then with 20% sucrose in PBS overnight at 4°C with agitation. Embryos were then incubated with 30% sucrose in PBS for 4 hours at room temperature, and then with a 1:1 mixture of Tissue-Tek O.C.T. Compound (O.C.T, Sakura, #4583) and 30% sucrose solution for 1 hour at room temperature with agitation. Finally, embryos were placed into 100% O.C.T for 10-20 minutes without agitation. Embryos were then positioned in fresh 100% OCT solution in plastic molds and frozen by dipping the mold into 2-methyl butane chilled on dry ice. 8 µm-thick sections were obtained using a Leica CM1950 Cryostat. For long term storage, slides were kept at −80°C. For immunofluorescence, we used an ImmEdge Hydrophobic Pen (VWR, 101098-065) to draw a hydrophobic barrier around tissue sections. Tissue sections were permeabilized by washing with PBS containing 0.05% Tween-20 (Sigma, cat # P7949) 3 times for 5 minutes. Then, slides were placed in a humidified chamber, and sections were incubated with a blocking buffer containing 5% donkey serum in PBS-0.05% Tween for 30 minutes at room temperature. The blocking buffer was then removed and replaced with primary antibodies diluted in blocking buffer and incubated with sections overnight at 4°C. After washing the slides 3 times for 5 minutes with PBS-0.05% Tween; sections were incubated with secondary antibodies for 1 hour at room temperature. Finally, slices were washed 3 times for 5 minutes each with PBS-0.05%Tween and mounted using a 50% v/v Glycerol, 50% v/v Methanol solution. Sections stained with SiR-Actin (Cytoskeleton, #CY-SC001, 2μM), samples were mounted with Vectashield Antifade Mounting Medium (Vector laboratories, #H-1000).

### TUNEL Assay

TUNEL assay was performed using the In Situ Cell Death Detection Kit from Roche (cat #11684795910) following the manufacturer’s instructions.

### Imaging, Quantifications, and Statistical Analyses

Embryos were imaged using confocal microscopy using Nikon A1R microscope with 20x CFI Apo LWD Lambda S water immersion objective (#MRD77200) or 25x CFI Plan Apo Lambda S silicone oil objectives (#MRD73250). Cultured cells were imaged using confocal microscopy and the Plan Fluor 567 40x Oil immersion objective (numerical aperture 1.3, # MRH01401). 3D reconstructions, surfacing, and quantifications were performed using IMARIS software (Bitplane, USA). Fluorescence intensity and angles were measured using Fiji software and plotted using Orient software, version 3.1.1. Cell ellipticity was measured using Imaris as described (Andrews et al., 2021). Statistical analyses were performed using Prism 9, version 9.4.1 (GraphPad Software, LLC).

## Acknowledgments

We thank Richard Hynes for providing *Fn1^+/-^* mice, Heiko Lickert for *Sox17^2A-iCre/+^* mice, Yumiko Saga for providing *Mesp1^Cre/+^* mice, and Reinhard Fässler for providing *Fn1^flox/flox^* mice. We thank Sam Russo for help with mouse colony maintenance and genotyping, and members of the Astrof lab for careful reading of the manuscript and helpful comments and suggestions throughout this work.

## Literature Cited

Alfano, D., A. Altomonte, C. Cortes, M. Bilio, R.G. Kelly, and A. Baldini. 2019. Tbx1 regulates extracellular matrix-cell interactions in the second heart field. Hum Mol Genet. 28:2295–2308.

Anderson, J.M., J. Li, and T.A. Springer. 2022. Regulation of integrin alpha5beta1 conformational states and intrinsic affinities by metal ions and the ADMIDAS. Mol Biol Cell. 33:ar56.

Andrews, T.G.R., W. Ponisch, E.K. Paluch, B.J. Steventon, and E. Benito-Gutierrez. 2021. Single-cell morphometrics reveals ancestral principles of notochord development. Development. 148.

Arriagada, C., V.A. Cavieres, C. Luchsinger, A.E. Gonzalez, V.C. Munoz, J. Cancino, P.V. Burgos, and G.A. Mardones. 2020. GOLPH3 Regulates EGFR in T98G Glioblastoma Cells by Modulating Its Glycosylation and Ubiquitylation. Int J Mol Sci. 21.

Aureille, J., V. Buffiere-Ribot, B.E. Harvey, C. Boyault, L. Pernet, T. Andersen, G. Bacola, M. Balland, S. Fraboulet, L. Van Landeghem, and C. Guilluy. 2019. Nuclear envelope deformation controls cell cycle progression in response to mechanical force. EMBO Rep. 20:e48084.

Avnur, Z., and B. Geiger. 1981. The removal of extracellular fibronectin from areas of cell-substrate contact. Cell. 25:121–132.

Bajolle, F., S. Zaffran, R.G. Kelly, J. Hadchouel, D. Bonnet, N.A. Brown, and M.E. Buckingham. 2006. Rotation of the myocardial wall of the outflow tract is implicated in the normal positioning of the great arteries. Circ Res. 98:421–428.

Bajolle, F., S. Zaffran, S.M. Meilhac, M. Dandonneau, T. Chang, R.G. Kelly, and M.E. Buckingham. 2008. Myocardium at the base of the aorta and pulmonary trunk is prefigured in the outflow tract of the heart and in subdomains of the second heart field. Dev Biol. 313:25–34.

Bohmer, R.M., E. Scharf, and R.K. Assoian. 1996. Cytoskeletal integrity is required throughout the mitogen stimulation phase of the cell cycle and mediates the anchorage-dependent expression of cyclin D1. Mol Biol Cell. 7:101–111.

Borreguero-Munoz, N., G.C. Fletcher, M. Aguilar-Aragon, A. Elbediwy, Z.I. Vincent-Mistiaen, and B.J. Thompson. 2019. The Hippo pathway integrates PI3K-Akt signals with mechanical and polarity cues to control tissue growth. PLoS Biol. 17:e3000509.

Cai, C.L., X. Liang, Y. Shi, P.H. Chu, S.L. Pfaff, J. Chen, and S. Evans. 2003. Isl1 identifies a cardiac progenitor population that proliferates prior to differentiation and contributes a majority of cells to the heart. Dev Cell. 5:877–889.

Chen, D., X. Wang, D. Liang, J. Gordon, A. Mittal, N. Manley, K. Degenhardt, and S. Astrof. 2015. Fibronectin signals through integrin alpha5beta1 to regulate cardiovascular development in a cell type-specific manner. Dev Biol. 407:195–210.

Chiang, H.Y., V.A. Korshunov, A. Serour, F. Shi, and J. Sottile. 2009. Fibronectin is an important regulator of flow-induced vascular remodeling. Arterioscler Thromb Vasc Biol. 29:1074–1079.

Chiquet-Ehrismann, R., P. Kalla, C.A. Pearson, K. Beck, and M. Chiquet. 1988. Tenascin interferes with fibronectin action. Cell. 53:383–390.

Chiquet-Ehrismann, R., and R.P. Tucker. 2011. Tenascins and the importance of adhesion modulation. Cold Spring Harb Perspect Biol. 3.

Clark, K., R. Pankov, M.A. Travis, J.A. Askari, A.P. Mould, S.E. Craig, P. Newham, K.M. Yamada, and M.J. Humphries. 2005. A specific alpha5beta1-integrin conformation promotes directional integrin translocation and fibronectin matrix formation. J Cell Sci. 118:291–300.

Cortes, C., A. Francou, C. De Bono, and R.G. Kelly. 2018. Epithelial Properties of the Second Heart Field. Circ Res. 122:142–154.

Cseh, B., S. Fernandez-Sauze, D. Grall, S. Schaub, E. Doma, and E. Van Obberghen-Schilling. 2010. Autocrine fibronectin directs matrix assembly and crosstalk between cell-matrix and cell-cell adhesion in vascular endothelial cells. J Cell Sci. 123:3989–3999.

Cui, Y., F.M. Hameed, B. Yang, K. Lee, C.Q. Pan, S. Park, and M. Sheetz. 2015. Cyclic stretching of soft substrates induces spreading and growth. Nat Commun. 6:6333.

Deno, D.C., T.M. Saba, and E.P. Lewis. 1983. Kinetics of endogenously labeled plasma fibronectin: incorporation into tissues. Am J Physiol. 245:R564–575.

Dupont, S. 2016. Role of YAP/TAZ in cell-matrix adhesion-mediated signalling and mechanotransduction. Exp Cell Res. 343:42–53.

Dupont, S., L. Morsut, M. Aragona, E. Enzo, S. Giulitti, M. Cordenonsi, F. Zanconato, J. Le Digabel, M. Forcato, S. Bicciato, N. Elvassore, and S. Piccolo. 2011. Role of YAP/TAZ in mechanotransduction. Nature. 474:179–183.

Dupont, S., and S.A. Wickstrom. 2022. Mechanical regulation of chromatin and transcription. Nat Rev Genet. 23:624–643.

Dyer, L.A., and M.L. Kirby. 2009. The role of secondary heart field in cardiac development. Dev Biol. 336:137–144.

Dyer, L.A., F.A. Makadia, A. Scott, K. Pegram, M.R. Hutson, and M.L. Kirby. 2010. BMP signaling modulates hedgehog-induced secondary heart field proliferation. Dev Biol. 348:167–176.

Dzamba, B.J., K.R. Jakab, M. Marsden, M.A. Schwartz, and D.W. DeSimone. 2009. Cadherin adhesion, tissue tension, and noncanonical Wnt signaling regulate fibronectin matrix organization. Dev Cell. 16:421–432.

Elbediwy, A., Z.I. Vincent-Mistiaen, B. Spencer-Dene, R.K. Stone, S. Boeing, S.K. Wculek, J. Cordero, E.H. Tan, R. Ridgway, V.G. Brunton, E. Sahai, H. Gerhardt, A. Behrens, I. Malanchi, O.J. Sansom, and B.J. Thompson. 2016. Integrin signalling regulates YAP and TAZ to control skin homeostasis. Development. 143:1674–1687.

Elosegui-Artola, A., I. Andreu, A.E.M. Beedle, A. Lezamiz, M. Uroz, A.J. Kosmalska, R. Oria, J.Z. Kechagia, P. Rico-Lastres, A.L. Le Roux, C.M. Shanahan, X. Trepat, D. Navajas, S. Garcia-Manyes, and P. Roca-Cusachs. 2017. Force Triggers YAP Nuclear Entry by Regulating Transport across Nuclear Pores. Cell. 171:1397–1410 e1314.

Engert, S., W.P. Liao, I. Burtscher, and H. Lickert. 2009. Sox17-2A-iCre: a knock-in mouse line expressing Cre recombinase in endoderm and vascular endothelial cells. Genesis. 47:603–610.

Engleka, K.A., L.J. Manderfield, R.D. Brust, L. Li, A. Cohen, S.M. Dymecki, and J.A. Epstein. 2012. Islet1 derivatives in the heart are of both neural crest and second heart field origin. Circ Res. 110:922–926.

Francou, A., C. De Bono, and R.G. Kelly. 2017. Epithelial tension in the second heart field promotes mouse heart tube elongation. Nat Commun. 8:14770.

Francou, A., E. Saint-Michel, K. Mesbah, and R.G. Kelly. 2014. TBX1 regulates epithelial polarity and dynamic basal filopodia in the second heart field. Development. 141:4320–4331.

Frank, D.U., L.K. Fotheringham, J.A. Brewer, L.J. Muglia, M. Tristani-Firouzi, M.R. Capecchi, and A.M. Moon. 2002. An Fgf8 mouse mutant phenocopies human 22q11 deletion syndrome. Development. 129:4591–4603.

Friedl, P., and R. Mayor. 2017. Tuning Collective Cell Migration by Cell-Cell Junction Regulation. Cold Spring Harb Perspect Biol. 9.

Galante, L.L., and J.E. Schwarzbauer. 2007. Requirements for sulfate transport and the diastrophic dysplasia sulfate transporter in fibronectin matrix assembly. J Cell Biol. 179:999–1009.

Garavito-Aguilar, Z.V., H.E. Riley, and D. Yelon. 2010. Hand2 ensures an appropriate environment for cardiac fusion by limiting Fibronectin function. Development. 137:3215–3220.

George, E.L., H.S. Baldwin, and R.O. Hynes. 1997. Fibronectins are essential for heart and blood vessel morphogenesis but are dispensable for initial specification of precursor cells. Blood. 90:3073–3081.

George, E.L., E.N. Georges-Labouesse, R.S. Patel-King, H. Rayburn, and R.O. Hynes. 1993. Defects in mesoderm, neural tube and vascular development in mouse embryos lacking fibronectin. Development. 119:1079–1091.

Giancotti, F.G., and G. Tarone. 2003. Positional control of cell fate through joint integrin/receptor protein kinase signaling. Annu Rev Cell Dev Biol. 19:173–206.

Hirsh, G.D., B.J. Dzamba, P.R. Sonavane, D.R. Shook, C.M. Allen, and D.W. DeSimone. 2018. Multiple functions for the catenin family member plakoglobin in cadherin-dependent adhesion, fibronectin matrix assembly, and *Xenopus* gastrulation movements. bioRxiv:318774.

Huang, W., R. Chiquet-Ehrismann, J.V. Moyano, A. Garcia-Pardo, and G. Orend. 2001. Interference of tenascin-C with syndecan-4 binding to fibronectin blocks cell adhesion and stimulates tumor cell proliferation. Cancer Res. 61:8586–8594.

Humphries, J.D., A. Byron, and M.J. Humphries. 2006. Integrin ligands at a glance. J Cell Sci. 119:3901–3903.

Hutson, M.R., X.L. Zeng, A.J. Kim, E. Antoon, S. Harward, and M.L. Kirby. 2010. Arterial pole progenitors interpret opposing FGF/BMP signals to proliferate or differentiate. Development. 137:3001–3011.

Hutson, M.R., P. Zhang, H.A. Stadt, A.K. Sato, Y.X. Li, J. Burch, T.L. Creazzo, and M.L. Kirby. 2006. Cardiac arterial pole alignment is sensitive to FGF8 signaling in the pharynx. Developmental biology. 295:486–497.

Hynes, R.O. 2002. Integrins: bidirectional, allosteric signaling machines. Cell. 110:673–687.

Hynes, R.O., E.L. George, E.N. Georges, J.L. Guan, H. Rayburn, and J.T. Yang. 1992. Toward a genetic analysis of cell-matrix adhesion. Cold Spring Harb Symp Quant Biol. 57:249–258.

Ilagan, R., R. Abu-Issa, D. Brown, Y.P. Yang, K. Jiao, R.J. Schwartz, J. Klingensmith, and E.N. Meyers. 2006. Fgf8 is required for anterior heart field development. Development. 133:2435–2445.

Imanaka-Yoshida, K., K. Matsumoto, M. Hara, T. Sakakura, and T. Yoshida. 2003. The dynamic expression of tenascin-C and tenascin-X during early heart development in the mouse. Differentiation. 71:291–298.

Jerome, L.A., and V.E. Papaioannou. 2001. DiGeorge syndrome phenotype in mice mutant for the T-box gene, Tbx1. *Nat Genet*. 27:286-291.

Kelly, R.G. 2023. The heart field transcriptional landscape at single-cell resolution. Dev Cell. 58:257–266.

Kelly, R.G., N.A. Brown, and M.E. Buckingham. 2001. The arterial pole of the mouse heart forms from Fgf10-expressing cells in pharyngeal mesoderm. Dev Cell. 1:435–440.

Kim, N.G., and B.M. Gumbiner. 2015. Adhesion to fibronectin regulates Hippo signaling via the FAK-Src-PI3K pathway. J Cell Biol. 210:503–515.

Kirby, M.L. 2007. Cardiac Development. Oxford University Press, New York.

Klass, C.M., J.R. Couchman, and A. Woods. 2000. Control of extracellular matrix assembly by syndecan-2 proteoglycan. J Cell Sci. 113 (Pt 3):493–506.

Kraft-Sheleg, O., S. Zaffryar-Eilot, O. Genin, W. Yaseen, S. Soueid-Baumgarten, O. Kessler, T. Smolkin, G. Akiri, G. Neufeld, Y. Cinnamon, and P. Hasson. 2016. Localized LoxL3-Dependent Fibronectin Oxidation Regulates Myofiber Stretch and Integrin-Mediated Adhesion. Dev Cell. 36:550–561.

Lackner, S., J. Schwendinger-Schreck, D. Julich, and S.A. Holley. 2013. Segmental assembly of fibronectin matrix requires rap1b and integrin alpha5. Dev Dyn. 242:122–131.

Li, D., A. Angermeier, and J. Wang. 2019. Planar cell polarity signaling regulates polarized second heart field morphogenesis to promote both arterial and venous pole septation. Development. 146.

Li, D., T. Sinha, R. Ajima, H.S. Seo, T.P. Yamaguchi, and J. Wang. 2016. Spatial regulation of cell cohesion by Wnt5a during second heart field progenitor deployment. Dev Biol. 412:18–31.

Li, D., and J. Wang. 2018. Planar Cell Polarity Signaling in Mammalian Cardiac Morphogenesis. Pediatr Cardiol. 39:1052–1062.

Liang, D., X. Wang, A. Mittal, S. Dhiman, S.Y. Hou, K. Degenhardt, and S. Astrof. 2014. Mesodermal expression of integrin alpha5beta1 regulates neural crest development and cardiovascular morphogenesis. Dev Biol. 395:232–244.

Lindsay, E., A. Botta, J. Vesna, S. Carattini-Rivera, Y.-C. Cheah, H.M. Rosenblatt, A. Bradley, and A. Baldini. 1999. Congenital heart disease in mice deficient for the DiGeorge syndrome region. Nature. 401:379–383.

Lindsay, E.A., F. Vitelli, H. Su, M. Morishima, T. Huynh, T. Pramparo, V. Jurecic, G. Ogunrinu, H.F. Sutherland, P.J. Scambler, A. Bradley, and A. Baldini. 2001. Tbx1 haploinsufficieny in the DiGeorge syndrome region causes aortic arch defects in mice. Nature. 410:97–101.

Lu, J., A.D. Doyle, Y. Shinsato, S. Wang, M.A. Bodendorfer, M. Zheng, and K.M. Yamada. 2020. Basement Membrane Regulates Fibronectin Organization Using Sliding Focal Adhesions Driven by a Contractile Winch. Dev Cell. 52:631–646 e634.

Marsden, M., and D.W. DeSimone. 2001. Regulation of cell polarity, radial intercalation and epiboly in Xenopus: novel roles for integrin and fibronectin. Development. 128:3635–3647.

Marsden, M., and D.W. DeSimone. 2003. Integrin-ECM interactions regulate cadherin-dependent cell adhesion and are required for convergent extension in Xenopus. Curr Biol. 13:1182–1191.

Masumoto, A., and M.E. Hemler. 1993. Multiple activation states of VLA-4. Mechanistic differences between adhesion to CS1/fibronectin and to vascular cell adhesion molecule-1. J Biol Chem. 268:228–234.

Meilhac, S.M., and M.E. Buckingham. 2018. The deployment of cell lineages that form the mammalian heart. Nat Rev Cardiol. 15:705–724.

Merscher, S., B. Funke, J.A. Epstein, J. Heyer, A. Puech, M.M. Lu, R.J. Xavier, M.B. Demay, R.G. Russell, S. Factor, K. Tokooya, B.S. Jore, M. Lopez, R.K. Pandita, M. Lia, D. Carrion, H. Xu, H. Schorle, J.B. Kobler, P. Scambler, A. Wynshaw-Boris, A.I. Skoultchi, B.E. Morrow, and R. Kucherlapati. 2001. TBX1 is responsible for cardiovascular defects in velo-cardio-facial/DiGeorge syndrome. In Cell. Vol. 104. 619–629.

Midwood, K.S., and J.E. Schwarzbauer. 2002. Tenascin-C modulates matrix contraction via focal adhesion kinase- and Rho-mediated signaling pathways. Mol Biol Cell. 13:3601–3613.

Midwood, K.S., L.V. Valenick, H.C. Hsia, and J.E. Schwarzbauer. 2004. Coregulation of fibronectin signaling and matrix contraction by tenascin-C and syndecan-4. Mol Biol Cell. 15:5670–5677.

Mittal, A., M. Pulina, S.Y. Hou, and S. Astrof. 2010. Fibronectin and integrin alpha 5 play essential roles in the development of the cardiac neural crest. Mech Dev. 127:472–484.

Mittal, A., M. Pulina, S.Y. Hou, and S. Astrof. 2013. Fibronectin and integrin alpha 5 play requisite roles in cardiac morphogenesis. Dev Biol. 381:73–82.

Miyamoto, S., H. Teramoto, J.S. Gutkind, and K.M. Yamada. 1996. Integrins can collaborate with growth factors for phosphorylation of receptor tyrosine kinases and MAP kinase activation: roles of integrin aggregation and occupancy of receptors. J Cell Biol. 135:1633–1642.

Mjaatvedt, C.H., T. Nakaoka, R. Moreno-Rodriguez, R.A. Norris, M.J. Kern, C.A. Eisenberg, D. Turner, and R.R. Markwald. 2001. The outflow tract of the heart is recruited from a novel heart-forming field. Dev Biol. 238:97–109.

Muzumdar, M.D., B. Tasic, K. Miyamichi, L. Li, and L. Luo. 2007. A global double-fluorescent Cre reporter mouse. Genesis. 45:593–605.

Nelson, W.J. 2003. Adaptation of core mechanisms to generate cell polarity. Nature. 422:766–774.

Nomaru, H., Y. Liu, C. De Bono, D. Righelli, A. Cirino, W. Wang, H. Song, S.E. Racedo, A.G. Dantas, L. Zhang, C.L. Cai, C. Angelini, L. Christiaen, R.G. Kelly, A. Baldini, D. Zheng, and B.E. Morrow. 2021. Single cell multi-omic analysis identifies a Tbx1-dependent multilineage primed population in murine cardiopharyngeal mesoderm. Nat Commun. 12:6645.

Orend, G., W. Huang, M.A. Olayioye, N.E. Hynes, and R. Chiquet-Ehrismann. 2003. Tenascin-C blocks cell-cycle progression of anchorage-dependent fibroblasts on fibronectin through inhibition of syndecan-4. Oncogene. 22:3917–3926.

Pankov, R., E. Cukierman, B.Z. Katz, K. Matsumoto, D.C. Lin, S. Lin, C. Hahn, and K.M. Yamada. 2000. Integrin dynamics and matrix assembly: tensin-dependent translocation of alpha(5)beta(1) integrins promotes early fibronectin fibrillogenesis. J Cell Biol. 148:1075–1090.

Park, E.J., Y. Watanabe, G. Smyth, S. Miyagawa-Tomita, E. Meyers, J. Klingensmith, T. Camenisch, M. Buckingham, and A.M. Moon. 2008. An FGF autocrine loop initiated in second heart field mesoderm regulates morphogenesis at the arterial pole of the heart. Development. 135:3599–3610.

Perez Gonzalez, N. J. Tao, N.D. Rochman, D. Vig, E. Chiu, D. Wirtz, and S.X. Sun. 2018. Cell tension and mechanical regulation of cell volume. Mol Biol Cell. 29:0.

Peters, D.M., L.M. Portz, J. Fullenwider, and D.F. Mosher. 1990. Co-assembly of plasma and cellular fibronectins into fibrils in human fibroblast cultures. J Cell Biol. 111:249–256.

Pulina, M., D. Liang, and S. Astrof. 2014. Shape and position of the node and notochord along the bilateral plane of symmetry are regulated by cell-extracellular matrix interactions. Biol Open. 3:583–590.

Pulina, M.V., S.Y. Hou, A. Mittal, D. Julich, C.A. Whittaker, S.A. Holley, R.O. Hynes, and S. Astrof. 2011. Essential roles of fibronectin in the development of the left-right embryonic body plan. Dev Biol. 354:208–220.

Radwanska, A., D. Grall, S. Schaub, S.B.F. Divonne, D. Ciais, S. Rekima, T. Rupp, A. Sudaka, G. Orend, and E. Van Obberghen-Schilling. 2017. Counterbalancing anti-adhesive effects of Tenascin-C through fibronectin expression in endothelial cells. Sci Rep. 7:12762.

Ramirez, A., and S. Astrof. 2020. Visualization and Analysis of Pharyngeal Arch Arteries using Whole-mount Immunohistochemistry and 3D Reconstruction. J Vis Exp.

Rammah, M., M. Theveniau-Ruissy, R. Sturny, F. Rochais, and R.G. Kelly. 2022. PPARgamma and NOTCH Regulate Regional Identity in the Murine Cardiac Outflow Tract. Circ Res. 131:842–858.

Ramsbottom, S.A., V. Sharma, H.J. Rhee, L. Eley, H.M. Phillips, H.F. Rigby, C. Dean, B. Chaudhry, and D.J. Henderson. 2014. Vangl2-regulated polarisation of second heart field-derived cells is required for outflow tract lengthening during cardiac development. PLoS Genet. 10:e1004871.

Roovers, K., and R.K. Assoian. 2003. Effects of rho kinase and actin stress fibers on sustained extracellular signal-regulated kinase activity and activation of G(1) phase cyclin-dependent kinases. Mol Cell Biol. 23:4283–4294.

Rozario, T., B. Dzamba, G.F. Weber, L.A. Davidson, and D.W. DeSimone. 2009. The physical state of fibronectin matrix differentially regulates morphogenetic movements in vivo. Dev Biol. 327:386–398.

Saga, Y., N. Hata, S. Kobayashi, T. Magnuson, M.F. Seldin, and M.M. Taketo. 1996. MesP1: a novel basic helix-loop-helix protein expressed in the nascent mesodermal cells during mouse gastrulation. Development. 122:2769–2778.

Saga, Y., S. Miyagawa-Tomita, A. Takagi, S. Kitajima, J. Miyazaki, and T. Inoue. 1999. MesP1 is expressed in the heart precursor cells and required for the formation of a single heart tube. Development. 126:3437–3447.

Sakai, T., K.J. Johnson, M. Murozono, K. Sakai, M.A. Magnuson, T. Wieloch, T. Cronberg, A. Isshiki, H.P. Erickson, and R. Fassler. 2001. Plasma fibronectin supports neuronal survival and reduces brain injury following transient focal cerebral ischemia but is not essential for skin-wound healing and hemostasis. Nat Med. 7:324–330.

Saoncella, S., F. Echtermeyer, F. Denhez, J.K. Nowlen, D.F. Mosher, S.D. Robinson, R.O. Hynes, and P.F. Goetinck. 1999. Syndecan-4 signals cooperatively with integrins in a Rho-dependent manner in the assembly of focal adhesions and actin stress fibers. Proc Natl Acad Sci U S A. 96:2805–2810.

Sechler, J.L., H. Rao, A.M. Cumiskey, I. Vega-Colon, M.S. Smith, T. Murata, and J.E. Schwarzbauer. 2001. A novel fibronectin binding site required for fibronectin fibril growth during matrix assembly. J Cell Biol. 154:1081–1088.

Shi, F., and J. Sottile. 2011. MT1-MMP regulates the turnover and endocytosis of extracellular matrix fibronectin. J Cell Sci. 124:4039–4050.

Singh, P., and J.E. Schwarzbauer. 2014. Fibronectin matrix assembly is essential for cell condensation during chondrogenesis. J Cell Sci. 127:4420–4428.

Sinha, T., D. Li, M. Theveniau-Ruissy, M.R. Hutson, R.G. Kelly, and J. Wang. 2015. Loss of Wnt5a disrupts second heart field cell deployment and may contribute to OFT malformations in DiGeorge syndrome. Hum Mol Genet. 24:1704–1716.

Sinha, T., B. Wang, S. Evans, A. Wynshaw-Boris, and J. Wang. 2012. Disheveled mediated planar cell polarity signaling is required in the second heart field lineage for outflow tract morphogenesis. Dev Biol. 370:135–144.

Soh, B.S., K. Buac, H. Xu, E. Li, S.Y. Ng, H. Wu, J. Chmielowiec, X. Jiang, L. Bu, R.A. Li, C. Cowan, and K.R. Chien. 2014. N-cadherin prevents the premature differentiation of anterior heart field progenitors in the pharyngeal mesodermal microenvironment. Cell Res. 24:1420–1432.

Somlyo, A.P., and A.V. Somlyo. 1994. Signal transduction and regulation in smooth muscle. Nature. 372:231–236.

Stefanovic, S., B. Laforest, J.P. Desvignes, F. Lescroart, L. Argiro, C. Maurel-Zaffran, D. Salgado, E. Plaindoux, C. De Bono, K. Pazur, M. Theveniau-Ruissy, C. Beroud, M. Puceat, A. Gavalas, R.G. Kelly, and S. Zaffran. 2020. Hox-dependent coordination of mouse cardiac progenitor cell patterning and differentiation. Elife. 9.

Stein, C., A.F. Bardet, G. Roma, S. Bergling, I. Clay, A. Ruchti, C. Agarinis, T. Schmelzle, T. Bouwmeester, D. Schubeler, and A. Bauer. 2015. YAP1 Exerts Its Transcriptional Control via TEAD-Mediated Activation of Enhancers. PLoS Genet. 11:e1005465.

Stepp, M.A., W.P. Daley, A.M. Bernstein, S. Pal-Ghosh, G. Tadvalkar, A. Shashurin, S. Palsen, R.A. Jurjus, and M. Larsen. 2010. Syndecan-1 regulates cell migration and fibronectin fibril assembly. Exp Cell Res. 316:2322–2339.

Sun, Y., X. Liang, N. Najafi, M. Cass, L. Lin, C.L. Cai, J. Chen, and S.M. Evans. 2007. Islet 1 is expressed in distinct cardiovascular lineages, including pacemaker and coronary vascular cells. Dev Biol. 304:286–296.

Sun, Z., A. Schwenzer, T. Rupp, D. Murdamoothoo, R. Vegliante, O. Lefebvre, A. Klein, T. Hussenet, and G. Orend. 2018. Tenascin-C Promotes Tumor Cell Migration and Metastasis through Integrin alpha9beta1-Mediated YAP Inhibition. Cancer Res. 78:950–961.

Takagi, J., K. Strokovich, T.A. Springer, and T. Walz. 2003. Structure of integrin alpha5beta1 in complex with fibronectin. EMBO J. 22:4607–4615.

Takahashi, S., M. Leiss, M. Moser, T. Ohashi, T. Kitao, D. Heckmann, A. Pfeifer, H. Kessler, J. Takagi, H.P. Erickson, and R. Fassler. 2007. The RGD motif in fibronectin is essential for development but dispensable for fibril assembly. J Cell Biol. 178:167–178.

Tang, Y., R.G. Rowe, E.L. Botvinick, A. Kurup, A.J. Putnam, M. Seiki, V.M. Weaver, E.T. Keller, S. Goldstein, J. Dai, D. Begun, T. Saunders, and S.J. Weiss. 2013. MT1-MMP-dependent control of skeletal stem cell commitment via a beta1-integrin/YAP/TAZ signaling axis. Dev Cell. 25:402–416.

Theveniau-Ruissy, M., M. Dandonneau, K. Mesbah, O. Ghez, M.G. Mattei, L. Miquerol, and R.G. Kelly. 2008. The del22q11.2 candidate gene Tbx1 controls regional outflow tract identity and coronary artery patterning. Circ Res. 103:142–148.

Tomer, D., C. Arriagada, S. Munshi, B.E. Alexander, B. French, P. Vedula, V. Caorsi, A. House, M. Guvendiren, A. Kashina, J.E. Schwarzbauer, and S. Astrof. 2022. A new mechanism of fibronectin fibril assembly revealed by live imaging and super-resolution microscopy. J Cell Sci. 135.

Trinh, L.A., and D.Y. Stainier. 2004. Fibronectin regulates epithelial organization during myocardial migration in zebrafish. Dev Cell. 6:371–382.

Trinh, L.A., D. Yelon, and D.Y. Stainier. 2005. Hand2 regulates epithelial formation during myocardial diferentiation. Curr Biol. 15:441–446.

Tsao, C.W., A.W. Aday, Z.I. Almarzooq, A. Alonso, A.Z. Beaton, M.S. Bittencourt, A.K. Boehme, A.E. Buxton, A.P. Carson, Y. Commodore-Mensah, M.S.V. Elkind, K.R. Evenson, C. Eze-Nliam, J.F. Ferguson, G. Generoso, J.E. Ho, R. Kalani, S.S. Khan, B.M. Kissela, K.L. Knutson, D.A. Levine, T.T. Lewis, J. Liu, M.S. Loop, J. Ma, M.E. Mussolino, S.D. Navaneethan, A.M. Perak, R. Poudel, M. Rezk-Hanna, G.A. Roth, E.B. Schroeder, S.H. Shah, E.L. Thacker, L.B. VanWagner, S.S. Virani, J.H. Voecks, N.Y. Wang, K. Yaffe, S.S. Martin, E. American Heart Association Council on, C. Prevention Statistics, and S. Stroke Statistics. 2022. Heart Disease and Stroke Statistics-2022 Update: A Report From the American Heart Association. Circulation:CIR0000000000001052.

Turner, C.J., K. Badu-Nkansah, and R.O. Hynes. 2017. Endothelium-derived fibronectin regulates neonatal vascular morphogenesis in an autocrine fashion. Angiogenesis. 20:519–531.

van den Berg, G., R. Abu-Issa, B.A. de Boer, M.R. Hutson, P.A. de Boer, A.T. Soufan, J.M. Ruijter, M.L. Kirby, M.J. van den Hoff, and A.F. Moorman. 2009. A caudal proliferating growth center contributes to both poles of the forming heart tube. Circ Res. 104:179–188.

van der Flier, A., K. Badu-Nkansah, C.A. Whittaker, D. Crowley, R.T. Bronson, A. Lacy-Hulbert, and R.O. Hynes. 2010. Endothelial alpha5 and alphav integrins cooperate in remodeling of the vasculature during development. Development. 137:2439–2449.

Verzi, M.P., D.J. McCulley, S. De Val, E. Dodou, and B.L. Black. 2005. The right ventricle, outflow tract, and ventricular septum comprise a restricted expression domain within the secondary/anterior heart field. Dev Biol. 287:134–145.

Villegas, S.N., M. Rothova, M.E. Barrios-Llerena, M. Pulina, A.K. Hadjantonakis, T. Le Bihan, S. Astrof, and J.M. Brickman. 2013. PI3K/Akt1 signalling specifies foregut precursors by generating regionalized extra-cellular matrix. Elife. 2:e00806.

Wada, K., K. Itoga, T. Okano, S. Yonemura, and H. Sasaki. 2011. Hippo pathway regulation by cell morphology and stress fibers. Development. 138:3907–3914.

Waldo, K.L., M.R. Hutson, H.A. Stadt, M. Zdanowicz, J. Zdanowicz, and M.L. Kirby. 2005. Cardiac neural crest is necessary for normal addition of the myocardium to the arterial pole from the secondary heart field. Dev Biol. 281:66–77.

Waldo, K.L., D.H. Kumiski, K.T. Wallis, H.A. Stadt, M.R. Hutson, D.H. Platt, and M.L. Kirby. 2001. Conotruncal myocardium arises from a secondary heart field. Development. 128:3179–3188.

Walker, J.L., A.K. Fournier, and R.K. Assoian. 2005. Regulation of growth factor signaling and cell cycle progression by cell adhesion and adhesion-dependent changes in cellular tension. Cytokine Growth Factor Rev. 16:395–405.

Wang, X., and S. Astrof. 2016. Neural crest cell-autonomous roles of fibronectin in cardiovascular development. Development. 143:88–100.

Ward, C., H. Stadt, M. Hutson, and M.L. Kirby. 2005. Ablation of the secondary heart field leads to tetralogy of Fallot and pulmonary atresia. Dev Biol. 284:72–83.

Watanabe, Y., S. Miyagawa-Tomita, S.D. Vincent, R.G. Kelly, A.M. Moon, and M.E. Buckingham. 2010. Role of mesodermal FGF8 and FGF10 overlaps in the development of the arterial pole of the heart and pharyngeal arch arteries. Circ Res. 106:495–503.

Weber, G.F., M.A. Bjerke, and D.W. DeSimone. 2011. Integrins and cadherins join forces to form adhesive networks. J Cell Sci. 124:1183–1193.

Weber, G.F., M.A. Bjerke, and D.W. DeSimone. 2012. A mechanoresponsive cadherin-keratin complex directs polarized protrusive behavior and collective cell migration. Dev Cell. 22:104–115.

Wennerberg, K., L. Lohikangas, D. Gullberg, M. Pfaff, S. Johansson, and R. Fassler. 1996. Beta 1 integrin-dependent and -independent polymerization of fibronectin. J Cell Biol. 132:227–238.

Xia, M., W. Luo, H. Jin, and Z. Yang. 2019. HAND2-mediated epithelial maintenance and integrity in cardiac outflow tract morphogenesis. Development. 146.

Yang, J.T., B.L. Bader, J.A. Kreidberg, M. Ullman-Cullere, J.E. Trevithick, and R.O. Hynes. 1999. Overlapping and independent functions of fibronectin receptor integrins in early mesodermal development. Dev Biol. 215:264–277.

Yang, J.T., H. Rayburn, and R.O. Hynes. 1993. Embryonic mesodermal defects in alpha 5 integrin-deficient mice. Development. 119:1093–1105.

Yelbuz, T.M., M.A. Choma, L. Thrane, M.L. Kirby, and J.A. Izatt. 2002. Optical coherence tomography: a new high-resolution imaging technology to study cardiac development in chick embryos. Circulation. 106:2771–2774.

Zamir, E., M. Katz, Y. Posen, N. Erez, K.M. Yamada, B.Z. Katz, S. Lin, D.C. Lin, A. Bershadsky, Z. Kam, and B. Geiger. 2000. Dynamics and segregation of cell-matrix adhesions in cultured fibroblasts. Nat Cell Biol. 2:191–196.

Zhang, J., Y. Lin, Y. Zhang, Y. Lan, C. Lin, A.M. Moon, R.J. Schwartz, J.F. Martin, and F. Wang. 2008. Frs2alpha-deficiency in cardiac progenitors disrupts a subset of FGF signals required for outflow tract morphogenesis. Development. 135:3611–3622.

Zhang, Q., D. Carlin, F. Zhu, P. Cattaneo, T. Ideker, S.M. Evans, J. Bloomekatz, and N.C. Chi. 2021. Unveiling Complexity and Multipotentiality of Early Heart Fields. Circ Res. 129:474–487.

Zhou, X., R.G. Rowe, N. Hiraoka, J.P. George, D. Wirtz, D.F. Mosher, I. Virtanen, M.A. Chernousov, and S.J. Weiss. 2008. Fibronectin fibrillogenesis regulates three-dimensional neovessel formation. Genes Dev. 22:1231–1243.

