## Supplemental Figures for "The mesodermal source of fibronectin is required for heart morphogenesis and cardiac outflow tract elongation by regulating cell shape, polarity, and mechanotransduction in the second heart field"

**Figure S1**

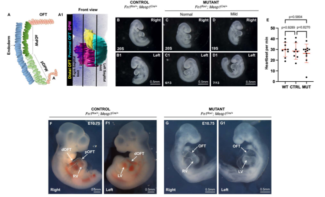


**Figure S1. $Fn1^{flox/-}$; $Mesp1^{Cre/+}$ mutant embryos do not survive past E10.75 of gestation. A.** Schematic representation of a sagittal optical section at the level displayed in the main manuscript figures; **A1.** Ventral view of E9.5 embryo with the left ventricle and the atrial chambers removed to show the DPW dorsal to the heart (pink). Analyses in this manuscript were performed within the DPW region between the left and right vertical lines, corresponding with the left and right lateral borders of the distal OFT. **B-D1.** Bright-field images of $Fn1^{flox/+}$;$Mesp1^{Cre/+}$ (Control) **B-B1,** and $Fn1^{flox/-}$;$Mesp1^{Cre/+}$ (Mutant) embryos, **C-C1, D-D1,** dissected at E9.5. **E.** Heartbeat per minute was measured in $Fn1^{flox/+}$; $Mesp1^{+/+}$ (WT), $Fn1^{flox/+}$;$Mesp1^{Cre/+}$ (Control) and $Fn1^{flox/-}$;$Mesp1^{Cre/+}$ (Mutant) embryos at E9.5 (20-26s). **F-F1.** $Fn1^{flox/+}$; $Mesp1^{Cre/+}$ (Control) and **G-G1.** $Fn1^{flox/-}$;$Mesp1^{Cre/+}$ (Mutant) embryos dissected at E10.75.

aDPW – anterior dorsal pericardial wall, pDWP- posterior dorsal pericardial wall, dOFT-distal outflow tract, pOFT-proximal outflow tract, A-atrium, RV – righr ventricle, LV-left ventricle. Dashed lines in **F** and **G** outline OFTs.

**Figure S2**

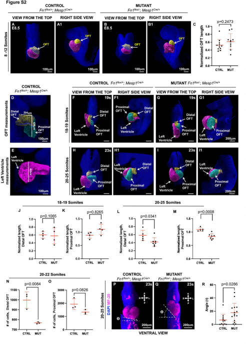


**Figure S2. Mesodermal Fn1 is required for OFT elongation at E9.5. A-A1. E8.5** $Fn1^{flox/+}$; $Mesp1^{Cre/+}$ (Control) and **B-B1.** $Fn1^{flox/-}$; $Mesp1^{Cre/+}$ (Mutant) embryos. Imaris was used to segment the OFT (yellow) and the left ventricle and atria (pink). **C.** The length of the OFT was normalized to the height of the left ventricle. N=11 control and 11 mutant embryos were analyzed. Each dot corresponds to one embryo.

**D-E**. Illustration of our method for measuring OFT dimensions. Imaris was used to segment the distal OFT (yellow), proximal OFT (turquoise), and the left ventricle and atria (pink). White lines show how the length of OFT segments and LV were measured. **F-I1. E9.5** $Fn1^{flox/+}$; $Mesp1^{Cre/+}$ (Control) and $Fn1^{flox/-}$; $Mesp1^{Cre/+}$ (Mutant) embryos were staged according to somite number. **F-G1.** Embryos with 18-19 somites. n=5 controls and 4 mutants. **H-I1.** Embryos with 20-25 somites, n= 5 control and 7 mutant embryos. All scale bars are 200 µm.

**J-M.** The length of the distal and proximal OFT in each embryo was normalized to the height of the left ventricle to control for embryo size.

**N-O**. Quantification of the number of myocardial cells in the distal and proximal OFT, n= 3 control and 3 mutant embryos. **P-Q.** $Fn1^{flox/+}$; $Mesp1^{Cre/+}$ (Control) and $Fn1^{flox/-}$; $Mesp1^{Cre/+}$ (Mutant) embryos were dissected at E9.5 (20-25s) and stained with DAPI and the MF-20 antibody. **R**. Quantification of angles in controls and mutants as shown in **P-Q**. n= 11 controls and 15 mutants. Means (horizontal bars) and standard deviations (error bars) are displayed; p was calculated using a 2-tailed, unpaired Student's t-test. OFT-outflow tract, MF-20- antibody detects sarcomeric myosin heavy chain.

Figure S3

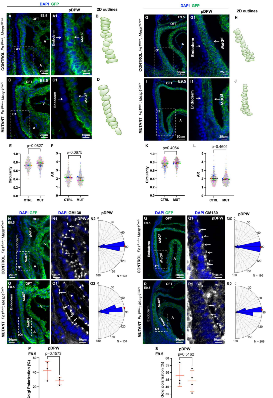


**Figure S3. Mesodermal Fn1 is not required for the regulation of the posterior DPW cell shape and orientation.** **A-B.** $Fn1^{flox/+}$; $Mesp1^{Cre/+}$ (Control) and **C-D.** $Fn1^{flox/-}$; $Mesp1^{Cre/+}$ (Mutant) embryos, dissected at E8.5 (9-11s) and stained to detect GFP (green) and nuclei (DAPI, blue). Regions marked by dashed rectangles in **A** and **C** are magnified in **A1** and **C1. B-D**. 2D-Outlines of pDPW cells. **E-F.** Shape analyses. Small dots mark data points from each cell; large dots mark the average per embryo. Data points from one embryo are marked in one color. 311 cells from 5 control and 326 cells from 5 mutant embryos were analyzed. **E.** Cell circularity. **F.** Aspect Ratio (AR). **G-H.** $Fn1^{flox/+}$; $Mesp1^{Cre/+}$ (Control) and **I-J.** $Fn1^{flox/-}$; $Mesp1^{Cre/+}$ (Mutant) embryos were dissected at E9.5 (21-25s) and stained to detect GFP (green) and nuclei (DAPI, blue). Regions marked by dashed rectangles in **G, I** are magnified in **G1-I1. H-J.** 2D-outlines of pDPW cells. **K-L.** Cell shape analyses. Small dots mark a data point from each cell; large dots mark the average per embryo. Data points from one embryo are marked with the same color; 390 cells from 5 control and 419 cells from 5 mutant embryos were analyzed. **K**. Cell circularity. **L.** Aspect Ratio (AR). Means (horizontal bars) and standard deviations (error bars) are displayed; p was calculated using a 2-tailed, unpaired Student's t-test using the means.

**N-N2.** $Fn1^{flox/+}$; $Mesp1^{Cre/+}$ (Control) and **O-O2.** $Fn1^{flox/-}$; $Mesp1^{Cre/+}$ (Mutant) embryos were dissected at E8.5 (9-11s), and stained to detect GFP (green), GM130 (white), and nuclei (DAPI, blue). **N1-O1.** Magnification of the pDPW. **N2-O2.** Semicircular histograms of the Golgi orientation angles in pDPW cells, measured as shown in **Fig. 2C. P.** Quantification of Golgi polarization. Golgi was considered polarized if the angle between the Golgi and the plane of the endoderm was between 85° and 95°. N=number of cells analyzed in 3 controls and 3 mutant embryos. **Q-Q2.** $Fn1^{flox/+}$; $Mesp1^{Cre/+}$ (Control) and **R-R2.** $Fn1^{flox/-}$; $Mesp1^{Cre/+}$ (Mutant) embryos were dissected at E9.5 (20-22s), and stained to detect GFP (green), GM130 (white), and nuclei (DAPI, blue). **Q1-R1.** Magnification of the pDPW. **Q2-R2.** Semicircular histograms of the Golgi orientation angles in pDPW cells, measured as above. **S.** Quantification of Golgi polarization. N=number of cells analyzed in 3 control and 4 mutant embryos were analyzed.

OFT-outflow tract, aDPW-anterior dorsal pericardial wall, pDPW-posterior dorsal pericardial wall, A-atrium.

Figure S4

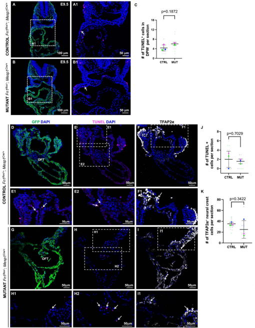


**Figure S4. Mesodermal Fn1 is not required for cell survival of DPW and neural crest cells at E9.5. A-A1.** Fn1$^{flox/+}$; Mesp1$^{Cre/+}$ (Control) and **B-B1.** Fn1$^{flox/-}$; Mesp1$^{Cre/+}$ (Mutant) embryos were dissected at E9.5 (21-26s), sectioned in the transverse orientation, stained to detect GFP (green), nuclei (DAPI, blue), and apoptosis using the TUNEL assay (magenta). Boxes in **A** and **B** are expanded in **A1** and **B1**. Arrows point to apoptotic cells (TUNEL$^+$) in the DPW. **C.** Quantification of TUNEL$^+$ cells in the DPW. 2-7 slices per embryo were analyzed. Small dots indicate the number of TUNEL$^+$ cells in one slice. Large dots mark the mean in each embryo. Data from the same embryo is displayed in the same color.

**D-F1.** Fn1$^{flox/+}$; Mesp1$^{Cre/+}$ (Control, n=3) and **G-I1.** Fn1$^{flox/-}$; Mesp1$^{Cre/+}$ (Mutant, n=3) embryos were dissected at E9.5 (21-26s), and transverse cryosections were used to perform TUNEL assay (magenta), Mesp1 lineage was detected by the expression of GFP (green), and neural crest cell (NCCs) were detected by the expression of TFAP2α (white), and nuclei were stained with DAPI, blue. Boxes in **E** and **H** are expanded in **E1, E2,** and **H1, H2,** where white arrows point to apoptotic cells (magenta) in the OFT. Boxes in **F** and **I** are expanded in **F1** and **I1,** where arrows point to NCCs (TFAP2α + cells). **J.** Quantification of TUNEL$^+$ cells per section. 2 - 7 sections per embryo were analyzed. Small dots in the plots mark the number of TUNEL$^+$ cells in each section. Large dots mark the mean in each embryo. **K.** Quantification of TFAP2α+ cells in the second pharyngeal arch. 2 – 7 sections per embryo were analyzed. Small dots mark the number of TFAP2α+ cells in each section.

Data from the same embryo are marked with the same color. Means (horizontal bars) and standard deviations (error bars) are displayed; Large dots mark the mean in each embryo. p values were calculated using a 2-tailed, unpaired Student's t-test using the means. DPW- dorsal pericardial wall, PHH3- phospho Histone H3, TUNEL- Terminal deoxynucleotidyl transferase (TdT) dUTP Nick-End Labeling.

**Figure S5**

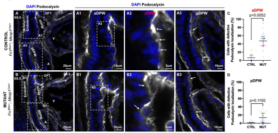


**Figure S5. A-D. Mesodermal Fn1 regulates podocalyxin localization in the anterior DPW at E8.5. A-A3.** $Fn1^{flox/+}$; $Mesp1^{Cre/+}$ (Control) and **B-B3.** $Fn1^{flox/-}$; $Mesp1^{Cre/+}$ (Mutant) embryos were dissected at E8.5 (9-11s) and stained to detect podocalyxin (white), and nuclei (DAPI, blue). Boxes in **A** and **B** were expanded in **A1-A2** and **B1-2** to show the anterior DPW and in **A3** and **B3** to show the posterior DPW. Arrows in **A2** point to cells with apical localization of podocalyxin. Arrows in **B2** point to the basolateral localization of podocalyxin in mutants. **A3, B3.** Podocalyxin is localized apically in the posterior DPW, arrows. **C-D.** Quantification of cells with defective podocalyxin localization in the anterior (**C**) and posterior DPW (**D**). 148 cells in the aDPW from 3 controls and 224 cells in pDPW from 3 mutants were analyzed. 3 controls and 3 mutants were analyzed.

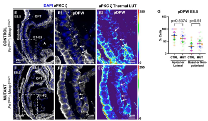


**E-E1. Mesodermal Fn1 is not required to regulate aPKC ζ localization in the posterior DPW at 8.5.** $Fn1^{flox/+}$; $Mesp1^{Cre/+}$ (Control) and **F-F1.** $Fn1^{flox/-}$; $Mesp1^{Cre/+}$ (Mutant) embryos (7-8 s) were stained to detect GFP (green), aPKC ζ (white), and nuclei (DAPI, blue). Boxes in **E** and **F** were expanded in **E1** and **F1,** and color-coded according to the signal intensity in **E2, F2.** Arrows show apical localization of aPKC ζ in control and mutant embryos. **G.** Quantification of cells with defective aPKC ζ localization in the anterior and posterior DPW. 3 controls and 3 mutants were analyzed.

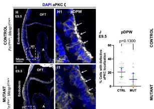


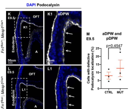


**H-M. Mesodermal Fn1 is not required for polarized expression of aPKC in the posterior DPW at E9.5.** Localization of aPKC ζ in the posterior DPW at E9.5 (20-22s). $Fn1^{flox/+}$; $Mesp1^{Cre/+}$ (Control, **H-H1**) and $Fn1^{flox/-}$; $Mesp1^{Cre/+}$ (Mutant, **I-I1**) embryos were stained for aPKC ζ (white) and nuclei (DAPI, blue). Boxes mark regions magnified to the right. Arrows in point aPKC ζ localization in control and mutant pDPW cells. **J.** Quantification of cells with defective aPKC ζ localization in the posterior DPW. 3 controls and 3 mutants were analyzed.

**K-M.** Localization of podocalyxin is apical in controls and mutants at E9.5 (19-24s). **M.** Quantification of cells in which podocalyxin was re-distributed away from the apical surface. Each dot marks mean number from one embryo.

**C, D, G, J.** Each small dot marks data from one slice, and each large dot is an average of all slices per embryo. Data from the same embryo is marked in one color. Means (horizontal lines) and standard deviations (error bars) are displayed; p was calculated using a 2-tailed, unpaired Student's t-test using means. A-atrium, aDPW-anterior dorsal pericardial wall, OFT-outflow tract, pDPW-posterior dorsal pericardial wall, V-ventricle.

Figure S6

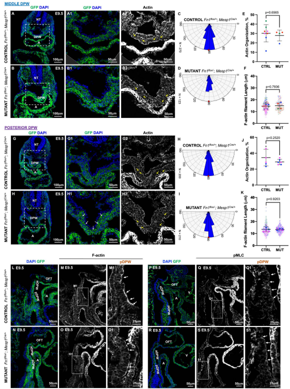


**Figure S6. The organization of the F-actin cytoskeleton in the middle and posterior DPW does not depend on the mesodermal Fn1. A-A2.** $Fn1^{flox/+}$; $Mesp1^{Cre/+}$ (Control) and **B-B2.** $Fn1^{flox/-}$; $Mesp1^{Cre/+}$ (Mutant) embryos were dissected at E9.5 (21-26s), and transverse cryosections from the middle of the DPW (see **Fig. 3E** for schematic) were stained to detect GFP (green), F-actin (white), and nuclei (DAPI, blue). Yellow arrows point to the F-actin cytoskeleton in the DPW. **C-D.** Quantification of F-actin angles in the middle DPW was measured relative to the apical surface. 270 cells from 4 controls and 123 cells from 3 mutants were analyzed. **E.** Percent of actin filaments oriented between 85° and 95° to the apical surface of the DPW. 270 cells from 4 controls and 123 cells from 3 mutants were analyzed. **F.** Quantification of F-actin length in the middle DPW cells.

**G-G2.** $Fn1^{flox/+}$; $Mesp1^{Cre/+}$ (Control) and **H-H2.** $Fn1^{flox/-}$; $Mesp1^{Cre/+}$ (Mutant) embryos were dissected at E9.5 (21-26s), and transverse cryosections from the posterior of the DPW (see **Fig. 3E** for schematic) were stained as in **A-B**. **H-I.** Quantification of F-actin angles in the posterior DPW was measured relative to the apical cell side. 200 cells from 3 controls and 210 cells from 2 mutants were analyzed as in **C-D**. **J.** Actin organization in cells of the posterior DPW was measured as in **E**. **K.** Quantification of F-actin length in the posterior DPW cells. Small dots mark data from each cell. Large dots are means from each optical slice. Data from the same embryo are displayed with the same color. Means (horizontal bars) and standard deviations (error bars) are displayed; p was calculated using a 2-tailed, unpaired Student's t-test, using means. DPW-dorsal pericardial wall, A- atrium, NT- Neural tube, OFT- outflow tract, V- ventricle

**L-M1.** $Fn1^{flox/+}$; $Mesp1^{Cre/+}$ (Control) and **N-O1.** $Fn1^{flox/-}$; $Mesp1^{Cre/+}$ (Mutant) embryos were dissected at E9.5 (21-26s), and sagittal cryosections were stained to detect GFP (green), F-actin (white) and nuclei (DAPI, blue). Boxes in **M and O** were expanded in **M1** and **O1** to show the posterior DPW. Arrows point to the apical enrichment of F-actin in the posterior of control and mutant embryos. **P-Q1.** $Fn1^{flox/+}$; $Mesp1^{Cre/+}$ (Control) and **R-S1.** $Fn1^{flox/-}$; $Mesp1^{Cre/+}$ (Mutant) embryos were dissected at E9.5 (21-26s), and sagittal cryosections were stained to detect GFP (green), pMLC (white) and nuclei (DAPI, blue). Boxes in **Q** and **S** are expanded in **Q1** and **S1** to show the posterior DPW at a higher magnification. Arrows point to cells with apical localization of pMLC.

Figure S7

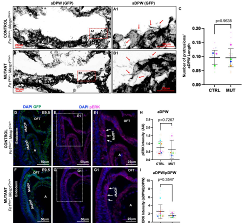


**Figure S7. Mesodermal depletion of Fn1 does not alter the number of basal protrusions or pErk1/2 levels at E9.5. A-A1.** Fn1$^{flox/+}$; Mesp1$^{Cre/+}$ (Control) and **B-B1.** Fn1$^{flox/-}$; Mesp1$^{Cre/+}$ (Mutant) embryos were dissected at E9.5 (21-26s), and transverse cryosections from the aDPW were stained to detect GFP (black). Red rectangles in **A, B** are expanded in **A1** and **B1**. Arrows point to protrusions extending from basal membranes of aDPW cells. **C.** The number of protrusions was normalized to the length of the aDPW. n= 3 control and n=4 mutant embryos. 2 – 7 sections per embryo were analyzed. Small dots mark the normalized number of basal protrusions in each section. Large dots mark the mean in each embryo. Data from the same embryo are displayed one color. **D-E1.** Fn1$^{flox/+}$; Mesp1$^{Cre/+}$ (Control) and **F-G1.** Fn1$^{flox/-}$; Mesp1$^{Cre/+}$ (Mutant) embryos were dissected at E9.5 (20-26s), fixed for 1 hour on ice, and stained to detect nuclei (DAPI, blue) and pERK (magenta). Regions outlined by dashed rectangles are expanded in **E1, G1** (anterior DPW). **H**. Quantification of the pERK intensity in the anterior DPW normalized by DAPI intensity. **I.** Quantification of pERK intensity ratio between anterior and posterior DPW. 5-7 optical slices from 6 controls and 4 mutant embryos were analyzed. Each small dot represents one slice, and each large dot is an average of all slices per embryo. Data from the same embryo are in one color. Means (horizontal bars) and standard deviations (error bars) are displayed, 2-tailed, unpaired Student's t-test was used to determine p values using means. aDPW-anterior dorsal pericardial wall, pDPW- posterior dorsal pericardial wall, OFT-outflow tract, A-atrium

**Figure S8**

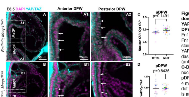


**Figure S8. Mesodermal depletion of Fn1 does not alter the nuclear translocation of YAP <u>at E8.5</u> in the DPW or in the <u>posterior</u> DPW at E9.5. A-D.** E8.5 (8-12s). **A-A2.** Fn1$^{flox/+}$; Mesp1$^{Cre/+}$ (Control) and **B-B2.** Fn1$^{flox/-}$; Mesp1$^{Cre/+}$ (Mutant) embryos were stained to detect nuclei (DAPI, magenta) and YAP protein (cyan). Regions outlined by dashed rectangles are expanded in **A1, B1** (anterior DPW) and **A2, B2** (posterior DPW). **C-D**. Quantification of the intensity ratio of nuclear and cytosolic YAP in the aDPW and pDPW. 5-6 optical slices from 2 controls and 4 mutant embryos were analyzed. Each small dot represents one slice, and each large dot is an average of all slices per embryo. Data from the same embryo is marked by the same color.

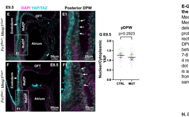


**E-G. E9.5 (23-26s), YAP protein staining in the <u>posterior</u> DPW**. **E-E1.** Fn1$^{flox/+}$; Mesp1$^{Cre/+}$ (Control) and **F-F1.** Fn1$^{flox/-}$; Mesp1$^{Cre/+}$ (Mutant) embryos were stained to detect nuclei (DAPI, magenta) and YAP protein (cyan). Regions outlined by dashed rectangles are expanded in **E1, F1** (posterior DPW). **G**. Quantification of the intensity ratio between nuclear and cytosolic YAP in pDPW. 7-8 optical slices from 4 controls and 7-8 from 4 mutant embryos were analyzed. Each small dot represents one slice, and each large dot is an average of all slices per embryo. Data from the same embryo are marked by the same color.

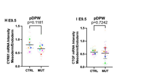


H. E9.5 (22-25s). **Quantification of *CYR61 mRNA* fluorescence intensity in the posterior DPW (mesoderm) normalized to the posterior endoderm**. n= 5 optical slices from 7 control and 6 mutant embryos were evaluated.

I. E9.5 (23-26s). **Quantification of *CTGF* mRNA fluorescence intensity in the posterior DPW (mesoderm) normalized to the posterior endoderm.** n= 6 optical slices from 4 control and 9 mutant embryos were evaluated. Each small dot represents one slice, and each large dot is an average of all slices per embryo. Data from the same embryo are marked by the same color. Means (horizontal bars) and standard deviations (error bars) are displayed, 2-tailed, unpaired Student's t-test was used to determine p values. aDPW-anterior dorsal pericardial wall, pDPW- posterior dorsal pericardial wall, OFT-outflow tract, A-atrium.

**Figure S9**

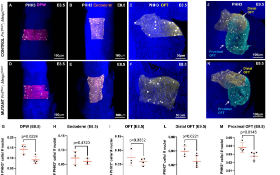


**Figure S9. Mesodermal Fn1 regulates DPW and OFT cell proliferation A-F.** $Fn1^{flox/+}$; $Mesp1^{Cre/+}$ (Control) and $Fn1^{flox/-}$; $Mesp1^{Cre/+}$ (Mutant) embryos at E8.5 (11-12s) were dissected and stained to detect PHH3 (white) and nuclei (DAPI, blue). **A, D.** DPW cells (magenta) and pHH3+ cells in the DPW (white) were segmented using Imaris. **B, E.** Endoderm (orange) and pHH3+ cells in the endoderm (white) were segmented using Imaris. **C, F.** Myocardial cells in the OFT were segmented in yellow, and the pHH3+ cells in the OFT were segmented in white using Imaris. **G**. Quantification of pHH3+ cells in the DPW of E8.5 embryos, n=3 control, and 4 mutant embryos. **H**. Quantification of pHH3+ cells in the endoderm of E8.5 embryos. n=3 for each genotype. **I**. Quantification of pHH3+ cells in the OFT of E8.5 embryos. n= 3 control and 4 mutant embryos.

**J-Z1.** $Fn1^{flox/+}$; $Mesp1^{Cre/+}$ (Control) and $Fn1^{flox/-}$; $Mesp1^{Cre/+}$ (Mutant) embryos at E9.5 (20-25s) were dissected and stained to detect pHH3 (white) and nuclei (DAPI, blue). **J, K.** Myocardial cells in the distal OFT were segmented in yellow, and proximal OFT – in turquoise, and the pHH3+ cells in each segment were marked in white using Imaris. **L-M**. Quantification of pHH3+ cells in the distal and proximal OFT myocardium, n=4 controls and 6 mutants.

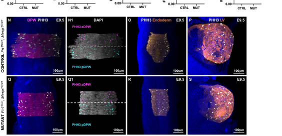


**N-N1, Q-Q1**. DPW (magenta) and pHH3+ cells in the DPW (white) were segmented using Imaris. pHH3+ cells in the aDPW are in magenta and pDPW are in turquoise. **O, R.** Endoderm (orange) and pHH3+ cells in the endoderm (white) were segmented using Imaris. **P, S.** Left ventricle (orange) and pHH3+ cells in the left ventricle (white) were segmented using Imaris.

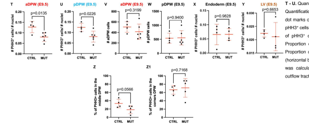


**T - U.** Quantification of pHH3+ cells in aDPW (**T**) and pDPW (**U**). **V-W**. Quantification of the number of cells in the aDPW and pDPW. Each dot marks one embryo, n=4 controls, 6 mutants. **X**. Quantification of pHH3+ cells in the endoderm. n=4 for each genotype **Y**. Quantification of pHH3+ cells in the left ventricle, n=4 for each genotype. **Z.** Proportion of pHH3+ cells in the medial region of the DPW. **Z1.** Proportion of pHH3+ cells at the lateral sides of the DPW. Means (horizontal bars) and standard deviations (error bars) are displayed; p was calculated using a 2-tailed, unpaired Student's t-test. OFT-outflow tract, pHH3 – phosphorylated Histone H3.

Figure S10

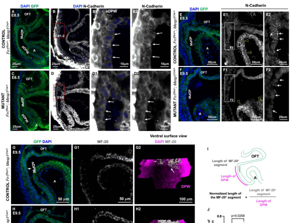


**Figure S10. Mesodermal Fn1 regulates the expression of N-Cadherin and suppresses precocious cardiac differentiation in the DPW. A-B2.** Fn1^flox/+^; Mesp1^Cre/+^ (Control) and **C-D2.** Fn1^flox/-^; Mesp1^Cre/+^ (Mutant) embryos were dissected at E8.5 (8-12s) and stained to detect GFP (green), N-cadherin (white) and nuclei (DAPI, blue). Boxes in **B-D** are expanded in **B1** and **D1.** Arrows point to aDPW cells. **E-E2.** Fn1^flox/+^; Mesp1^Cre/+^ (Control) and **F-F2.** Fn1^flox/-^; Mesp1^Cre/+^ (Mutant) embryos were dissected at E9.5 (21-26s), and stained to detect GFP (green), N-cadherin (white) , and nuclei (DAPI, blue). Boxed regions in **E1** and **F1** are expanded in **E2-F2**. Arrows point to junctional N-cadherin in controls (**E2**) and its absence in the mutants, **F2**. Yellow arrowheads point to junctional N-cadherin staining in the atrial myocardial cells in the same confocal plane.

**G-G2**. Fn1^flox/+^; Mesp1^Cre/+^ (Control) and **H-H2.** Fn1^flox/-^; Mesp1^Cre/+^ (Mutant) embryos were dissected at E9.5 (21-26s), and stained to detect GFP (green), sarcomeric myosin heavy chain (MF-20, white), and nuclei (DAPI, blue). The arrow in **H1** points to cells in the aDPW ectopically expressing sarcomeric myosin heavy chain. **G2, H2.** Ventral views of the DPW. 3D reconstructions of the DPW (pink) using Imaris. MF-20 staining in the DPW is marked by the white surface (arrows). **I**. Schematic representation of the method used to quantify MF-20 staining. **J.** The length of the MF20 staining relative to the DPW length in each section is plotted. Eight optical sections per embryo were analyzed from 5 controls and 8 mutants. Small dots mark data from each slice; Large dots mark means. Data from the same embryo are displayed with the same color. A 2-tailed, unpaired. Student's t-test was used to determine p values. DPW- dorsal pericardial wall, OFT- outflow tract, A- atrium. MF-20- sarcomeric myosin heavy chain.

**Figure S11**

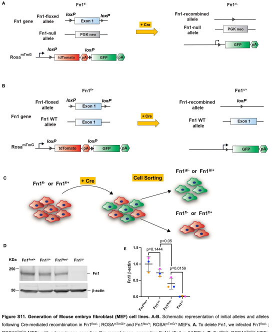


**Figure S11. Generation of Mouse embryo fibroblast (MEF) cell lines. A-B.** Schematic representation of initial alleles and alleles following Cre-mediated recombination in $Fn1^{flox/-}$; $ROSA^{mTmG/+}$ and $Fn1^{flox/+}$; $ROSA^{mTmG/+}$ MEFs. **A.** To delete Fn1, we infected $Fn1^{flox/-}$; $ROSA^{mTmG/+}$ MEFs with adenoviruses encoding Cre recombinase, generating $Fn1^{\Delta/-}$ (Fn1-null MEFs). **B.** $Fn1^{flox/+}$; $ROSA^{mTmG/+}$ MEFs were treated with adenoviruses encoding Cre recombinase to generate additional control MEFs with the genotype $Fn1^{\Delta/+}$. **C.** After infection, cells were expanded and sorted. Membrane-tagged GFP-expressing cells are in green and uninfected, membrane-tagged td-Tomato-expressing cells are in red. **D.** Western blot using anti-Fn1 antibody shows that we successfully ablated Fn1 in $Fn1^{\Delta/-}$ (Fn1-null) MEFs. β-actin was used as a loading control. **E.** Densitometrical quantification of Fn1 levels relative to β-actin. Means of three independent experiments (horizontal bars) and standard deviations (error bars) are displayed; p was calculated using a 2-tailed, unpaired Student's t-test.

**Figure S12**

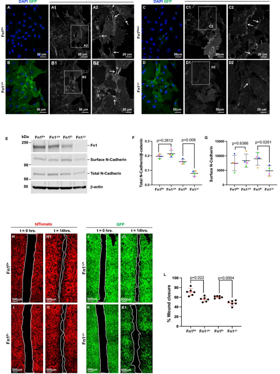


**Figure S12. Fn1 regulates N-Cadherin cell expression, localization at adherens junctions, and collective cell migration in *vitro*.**

**A-D2** Immunostaining to detect GFP (green), N-cadherin (white), and nuclei (DAPI, blue) in $Fn1^{flox/+}$, $Fn1^{\Delta/+}$, $Fn1^{flox/-}$, and $Fn1^{\Delta/-}$ (Fn1-null) MEFs. Regions marked by dashed rectangles in **A1-B1** and **C1-D1** are expanded in **A2-B2** and **C2-D2.** Arrows point to N-cadherin at cell borders in control MEFs (**A2, B2, and C2**) and to the absence of N-cadherin at cell-cell contacts in Fn1-null MEFs (**D2**).

**E-G.** Western blot analysis of total cell lysates and cell surface proteins showed the downregulation of total N-Cadherin levels in Fn1-null MEFs. **E.** Western blot. Positions of molecular weight markers are indicated on the left. **F-G.** Densitometrical quantification of immunoblot signals. N = 4 for each genotype. Each experiment is represented by a dot of a different color. Means (horizontal bars) and standard deviations (error bars) are displayed, and 2-tailed, unpaired Student's t-tests were used to determine p values.

**H-L. Fn1 regulates collective cell migration. H, I, J, K.** Cells imaged immediately after the scratch wound (time=0 hrs). **H1, I1, J1, K1.** Cells imaged 14 hours after the scratch (time=14 hrs). The area remaining cell-free at t=14 hours is outlined.

**L.** To quantify % wound closure, cell-free area at t=14($A_{t=14}$) was subtracted from cell-free area at t=0 ($A_{t=0}$), and the % wound closure was calculated as $100^*(A_{t=0} - A_{t=14})/ A_{t=0}$); N=6 independent experiments. Means (horizontal bars) and standard deviations (error bars) are displayed; p values were calculated using 2-tailed, unpaired Student's t-tests.

**Figure S13**

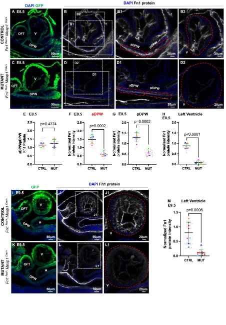


**Figure S13. Downregulation of Fn1 protein in the mesoderm of Fn1$^{flox/-}$; Mesp1$^{Cre/+}$ (Mutant) embryos at E8.5. A-H.** E8.5 embryos (8-12s). **A-B2.** Fn1$^{flox/+}$; Mesp1$^{Cre/+}$ (Control) and **C-D2.** Fn1$^{flox/-}$; Mesp1$^{Cre/+}$ (Mutant) embryos were stained to detect GFP (green), Fn1 protein (white), and nuclei (DAPI, blue).

**E.** Ratio of Fn1 protein intensity shows similar levels of Fn1 in the anterior and posterior DPW of control and mutant embryos. **F-H.** Fn1 protein levels decrease in the DPW and left ventricle of the mutants at E8.5. Fn1 intensity in the aDPW, pDWP, and left ventricle regions outlined by the red dash lines in **B1-2** and **D1-D2** was normalized by DAPI intensity; 5 optical slices per embryo from 5 control and 5 mutant embryos were evaluated.

**I-M.** E9.5 embryos (22-26s). **I-J1.**Fn1$^{flox/+}$; Mesp1$^{Cre/+}$ (Control) and **K-L1.** Fn1$^{flox/-}$; Mesp1$^{Cre/+}$ (Mutant) embryos were stained to detect Fn1 protein (white) and nuclei (DAPI, blue). Regions outlined by rectangles are expanded in **J1** and **L1**. **M**. Fn1 protein expression in the left ventricle. 7 optical slices per embryo from 8 control and 10 mutant embryos were evaluated. Each small dot represents one slice, and each large dot is an average of all slices per embryo. Data from the same embryo are marked by the same color. Means (horizontal bars) and standard deviations (error bars) are displayed; a 2-tailed, unpaired Student's t-test was used to determine p values using the means. aDPW-anterior dorsal pericardial wall, pDPW-posterior dorsal pericardial wall, OFT-outflow tract, A-atrium, LV-left ventricle.

**Figure S14**

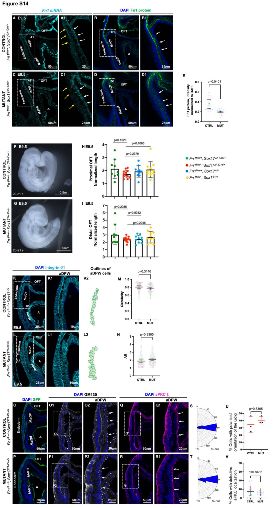


**Figure S14. Fn1 synthesis in the Sox17 lineage is not required for the regulation of cell shape or cell polarity in the DPW. A-B1** $Fn1^{flox/+}$; $Sox17^{2A-iCre/+}$ (Control) and **C-D1.** $Fn1^{flox/-}$; $Sox17^{2A-iCre/+}$ (Mutant) embryos were dissected at E9.5 (20-25s). **A** and **C. Expression of *Fn1 mRNA*.** Embryos were labeled with DAPI (blue) and anti-sense Fn1 probe (turquoise). Regions marked by the dashed rectangles in **A** and **C** are magnified in **A1** and **C1**. Yellow arrows point to the endoderm, white arrows point to the DPW mesoderm. **B** and **D. Expression of Fn1 protein**. Embryos were stained to detect Fn1 protein (green) and nuclei (DAPI, blue). Boxes in **B** and **D** were expanded in **B1** and **D1**. **E.** Quantification of Fn1 protein levels underlying the anterior DPW. Fn1 intensity in the aDPW was normalized by DAPI intensity; 5 optical slices per embryo from 3 control and 3 mutant embryos were evaluated. Each small dot represents one slice, and each large dot is an average of all slices per embryo. Data from the same embryo are marked by the same color. Means (horizontal bars) and standard deviations (error bars) are displayed, a 2-tailed, unpaired Student's t-test was used to determine the p - value using averages.

**F.** $Fn1^{flox/+}$; $Sox17^{2A-iCre/+}$ (Control) and **G.** $Fn1^{flox/-}$; $Sox17^{2A-iCre/+}$ (Mutant) embryos dissected at E9.5. **H-I** Quantification of the proximal and distal OFT lengths normalized by the height of the otic vesicle. Each dot marks one embryo. 2-tailed, unpaired Student's t-test was used to determine the p – values.

**K-K2.** $Fn1^{flox/+}$; $Sox17^{+/+}$ (Control) and **L-L2**. $Fn1^{flox/+}$; $Sox17^{2A-iCre/+}$ (Mutant) embryos were dissected at E9.5 (18-22s) and stained to detect integrin-β1 (turquoise) and nuclei (DAPI, blue). Regions marked by dashed rectangles in **K** and **L** are magnified in **K1-L1. K2-L2.** 2D-Outlines of aDPW cells. **M-N.** Cell shape analyses. **M**. Cell circularity **N.** AR (Aspect ratio). Color coding and statistical analyses were done as in panel **E**. 156 cells from 2 controls and 152 cells from 2 mutant embryos were analyzed.

**O-Q1.** $Fn1^{flox/+}$; $Sox17^{2A-iCre/+}$; $ROSA^{mTmG/+}$ (Control) and **P-R1.** $Fn1^{flox/-}$; $Sox17^{2A-iCre/+}$ $ROSA^{mTmG/+}$ (Mutant) embryos were dissected at E9.5 (21-23s) and stained to detect GFP (green), GM130 (white), aPKC ζ (pink), and nuclei (DAPI, blue). Regions marked by dashed rectangles are magnified in panels to the right. **S-T.** Semicircular histograms represent the distribution of the Golgi orientation angles in the aDPW, measured as in **Figure 2C**. N=total number of cells analyzed in 3 controls and 3 mutants. **U**. Quantification of Golgi polarization. Golgi was considered polarized if the angle between the Golgi and the plane of the endoderm fell between 85° and 95°. **V.** Quantification of cells in which aPKC ζ was re-distributed away from the apical surface. DPW cells from 5 different slices were evaluated in 3 control and 3 mutant embryos. Color coding and statistical analyses were done as in panel **E**. OFT-outflow tract, aDPW-anterior dorsal pericardial wall, pDPW-posterior dorsal pericardial wall, and A-atrium.

**Figure S15**

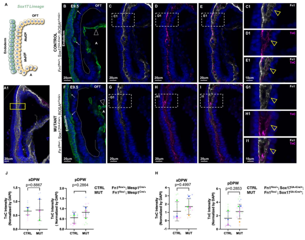


**Figure S15. The depletion of Fn1 in the Sox17 lineage does not alter TnC localization at the aDPW. A.** Schematic of the Cre expression in the Sox17 lineage, endodermal Sox17 lineage cells are marked in green. **A1.** A 20-micron-thick optical slice and an example of a region (yellow rectangle) where the distributions of Fn1 and TnC proteins were analyzed using Fiji in **Figure 7**. **B, F.** $Sox17^{2A-Cre/+}$ lineage labels the endoderm, endocardium (arrowheads), and endothelium (arrows) but not the DPW (dashed lines mark the apical surface of the DPW). **B-E1.** $Fn1^{flox/+}$; $Sox17^{2A-iCre/+}$; $ROSA^{mTmG/+}$ (Control) and **F-I1.** $Fn1^{flox/-}$; $Sox17^{2A-Cre/+}$; $ROSA^{mTmG/+}$ (Mutant) embryos were dissected at E9.5 (20-22s) and stained to detect GFP (green), Fn1 protein (white), TnC (magenta) and nuclei (DAPI, blue). Regions marked by dashed rectangles in **C-E** and **G-I** are magnified in **C1-E1** and **G1-I1**. Yellow arrowheads point to the co-localization of Fn1 and TnC. **J.** TnC protein levels are comparable in the DPW of $Fn1^{flox/+}$; $Mesp1^{Cre/+}$ controls and $Fn1^{flox/-}$; $Mesp1^{Cre/+}$ mutants. TnC staining intensity was normalized to DAPI intensity in each slice. 5 slices for each embryo were analyzed. **H.** TnC protein levels are comparable in the DPW of $Fn1^{flox/+}$; $Sox17^{2A-iCre/+}$ controls and $Fn1^{flox/-}$; $Sox17^{2A-iCre/+}$ endodermal mutants. TnC staining intensity was normalized to DAPI intensity in each slice. 5 slices per embryo were analyzed. Each small dot represents one slice, and each large dot is an average of all slices per embryo. Means (horizontal bars) and standard deviations (error bars) are displayed, 2-tailed, unpaired Student's t-test was used to determine p values using the means. OFT-outflow tract, A-atrium.
