## Supplemental Table 1 for "The mesodermal source of fibronectin is required for heart morphogenesis and cardiac outflow tract elongation by regulating cell shape, polarity, and mechanotransduction in the second heart field"

**Supplemental Table 1.** Genotypes of embryos isolated from crosses between Fn1^flox/flox^ and Fn1^+/-^; Mesp1^Cre/+^ animals.

| Genotype  Stage | FN1^flox/+^; Mesp1^+/+^ | FN1^flox/-^; Mesp1^+/+^ | FN1^flox/+^; Mesp1^Cre/+^ | FN1^flox/-^; Mesp1^Cre/+^ | Total number of embryos/pups genotyped |
| --- | --- | --- | --- | --- | --- |
| E9.5 | 12 | 10 | 13 | 12 | 47 |
| E10.75 | 24 | 20 | 14 | 7* | 65 |
| E13.5-E17.5 | 45 | 36 | 51 | 0 | 132 |

* Degenerating
